# Diurnal rhythmicity of fecal microbiota and metabolite profiles in the first year of life: a randomized controlled interventional trial with infant formula

**DOI:** 10.1101/2023.10.19.563092

**Authors:** Nina Heppner, Sandra Reitmeier, Marjolein Heddes, Michael Vig Merino, Leon Schwartz, Alexander Dietrich, Markus List, Michael Gigl, Chen Meng, Hélène Omer, Karin Kleigrewe, Melanie Schirmer, Daan R van der Veen, Silke Kiessling, Dirk Haller

## Abstract

Microbiota assembly in the infant gut is influenced by time and duration of dietary exposure to breast-milk, infant formula and solid foods. In this randomized controlled intervention study, longitudinal sampling of infant stool (n=998) showed similar development of fecal bacterial communities (16S rRNA/shallow metagenomics sequencing) between formula- and breast-fed infants during the first year of life (N=210). Infant formula supplemented with galacto-oligosaccharides (GOS) was most efficient to sustain high levels of bifidobacteria compared to formula containing *B. longum* and *B. breve* or placebo. In addition to primary endpoints, metabolite and bacterial profiling revealed 24-hour oscillations and data integration identified circadian networks. Rhythmicity in bacterial diversity, specific taxa and functional pathways increased with age and was strongest following breast-feeding and GOS-supplementation. Circadian rhythms in dominant taxa were discovered *ex-vivo* in a chemostat model. Hence, microbiota rhythmicity develops early in life, likely via bacterial intrinsic clock mechanism and is affected by diet.

## Main

Characteristic of the developing infant gut microbiota is the high degree of individual variation during the first years of life^1^. The deterministic or stochastic succession of microbial communities in the healthy infant gut is largely linked to environmental and dietary exposures^2–4^. Colonization of the sterile infant gut starts at birth and depends on fetal exposure to microbial metabolites from the mother (immune imprinting)^5^, birth mode, and the presence of microbial pioneers from the fecal, vaginal or skin environment^3^. Timing and duration of breast- and formula feeding are subsequent drivers of early life microbial succession^6,7^. Breast-fed infants are typically characterized by low bacterial diversity and dominance of bifidobacteria^8–10^. Human breastmilk oligosaccharides (HMOs) selectively shape the early life colonization^11–13^ and the supplementation of infant formula with complex oligosaccharides such as galacto-oligosaccharides (GOS) partially mimics the effect of breastmilk^14,15^. Considering the well-documented effects of the early life microbiota on health outcomes later in life^8,16,17^, the attempts to optimize infant formula are numerous^18–21^ but often lack well-controlled intervention strategies to directly compare supplements in infant formula.

Circadian clocks (lat. *circa* = approximately, *dies* = day) have developed in almost all organisms to anticipate recurring environmental changes. In mammals, a central clock in the brain orchestrates subordinated peripheral clocks of various tissues and single cells^22^. Thereby the circadian system drives 24-hour oscillations to control body and organ functions, however, the contribution of circadian rhythms in the developing gut including the process of microbial colonization is completely unknown^23^. So far, circadian clock mechanisms were believed to be restricted to photosynthetic bacteria, such as cyanobacteria, but, in single bacterial cultures circadian oscillations were found in gene expression of non-photosynthetic *Bacillus subtilis*^24^ and swarming behavior of the gut bacterium *Klebsiella aerogenes*^25^, suggesting the presence of endogenous circadian clocks in specific gut bacteria. In recent years, time-of-day dependent fluctuations in the abundance of certain gut bacterial species were shown at the individual and population level^26,27^. Growing evidence provides a link between loss of rhythmicity and disease pathology^27^. Animal studies determined the origin of these bacterial oscillations to reside in the circadian system^28–30^, particularly the intestinal circadian clock^31^. The mammalian circadian system develops *in utero,* which continues during aging and plays an important role during development^32^. Gut microbiota rhythms can influence clock gene expression in the host ^33,34^ and disturbance of the circadian crosstalk between bacteria and the host was shown to affect gastrointestinal health and metabolism^30,31,34^. Whether bacteria endogenous clock mechanisms can drive rhythmicity of complex microbiota and thereby impact host physiology remains unknown.

Here we established a randomized, controlled intervention trial including 210 infants to address the role of cow’s-milk based infant formula supplemented with bifidobacteria and GOS on early life development of the gut microbiota. Longitudinal sampling of infant stool (n=998) allowed us to temporally resolve microbiota and metabolite profiles in response to the supplemented infant formula. Exclusively breast-fed infants served as a reference group to evaluate the impact of infant formula feeding. In addition to the analysis of primary endpoints, we identified diurnal oscillations of the microbiome in relation to dietary exposure and confirmed the circadian rhythmicity of dominant taxa *ex-vivo* in a gut chemostat model.

## Results

Eligibility assessment of 265 pregnant women led to the inclusion of 223 infants in the intervention study. Of those infants, 210 actively participated and were randomly assigned to one of the four different formula groups (**Fig. 1**). 63 infants were breast-fed for the duration of the intervention (1 year). 35 infants were randomized to receive the placebo containing no additional supplements (Formula A), 39 infants were randomized to Formula B containing *B. longum* and *B. breve*, 36 infants received formula containing prebiotics (GOS) (Formula C) and 37 infants consumed formula containing both bifidobacteria and GOS (Formula D) (**Fig. 1**). Fecal samples were collected at five time points during the first year of life (**Fig. 2A**). 87% of participants contributed to all sampling time points. An additional, voluntary follow-up sample was provided by 93 participants at the 24-month time point (29 breast-fed, 15 formula A-fed, 16 formula B-fed, 17 formula C-fed, 16 formula D-fed infants). The characteristics of participants in the five study arms were comparable (**Table 1**) and the weight gain of infants was similar across groups (**Suppl. Fig. 1A**). Daily formula amounts consumed were similar across interventional groups (**Suppl. Fig. 1B-C**). Differences in breastmilk metabolites over 103 samples could not classify the corresponding infant fecal metabolite profiles (**Suppl. Fig. 1D-E**). Rates for study discontinuation were low (**Table 1**) with suspected cow’s milk protein allergy (6 out of 9 cases) being the main reason for study withdrawal.

**Figure 1.**
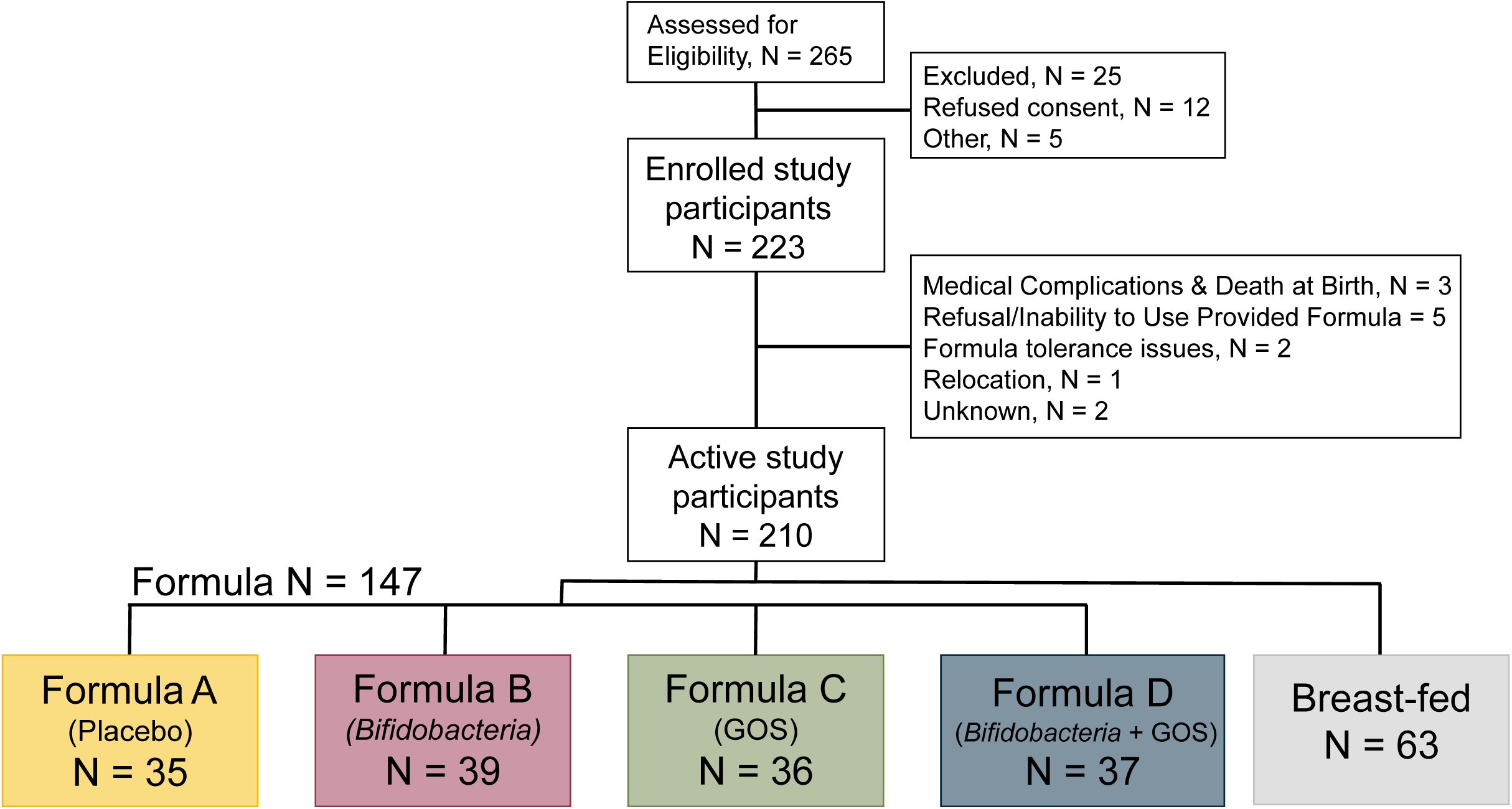
Study schematic of participant enrollment and randomization. Schematic of participant enrolment and reasons for drop-out before start of study. Active study participants (N =210), provided at least one fecal sample. Newborns were randomly assigned to one of four formula groups: Placebo, *Bifidobacterium*-supplemented (Formula B), galacto-oligosaccharide (GOS)-supplemented (Formula C) or formula containing GOS and *Bifidobacteria* (Formula D). Breast-fed infants that never consumed any of the provided formula served as reference group.

**Figure 2.**
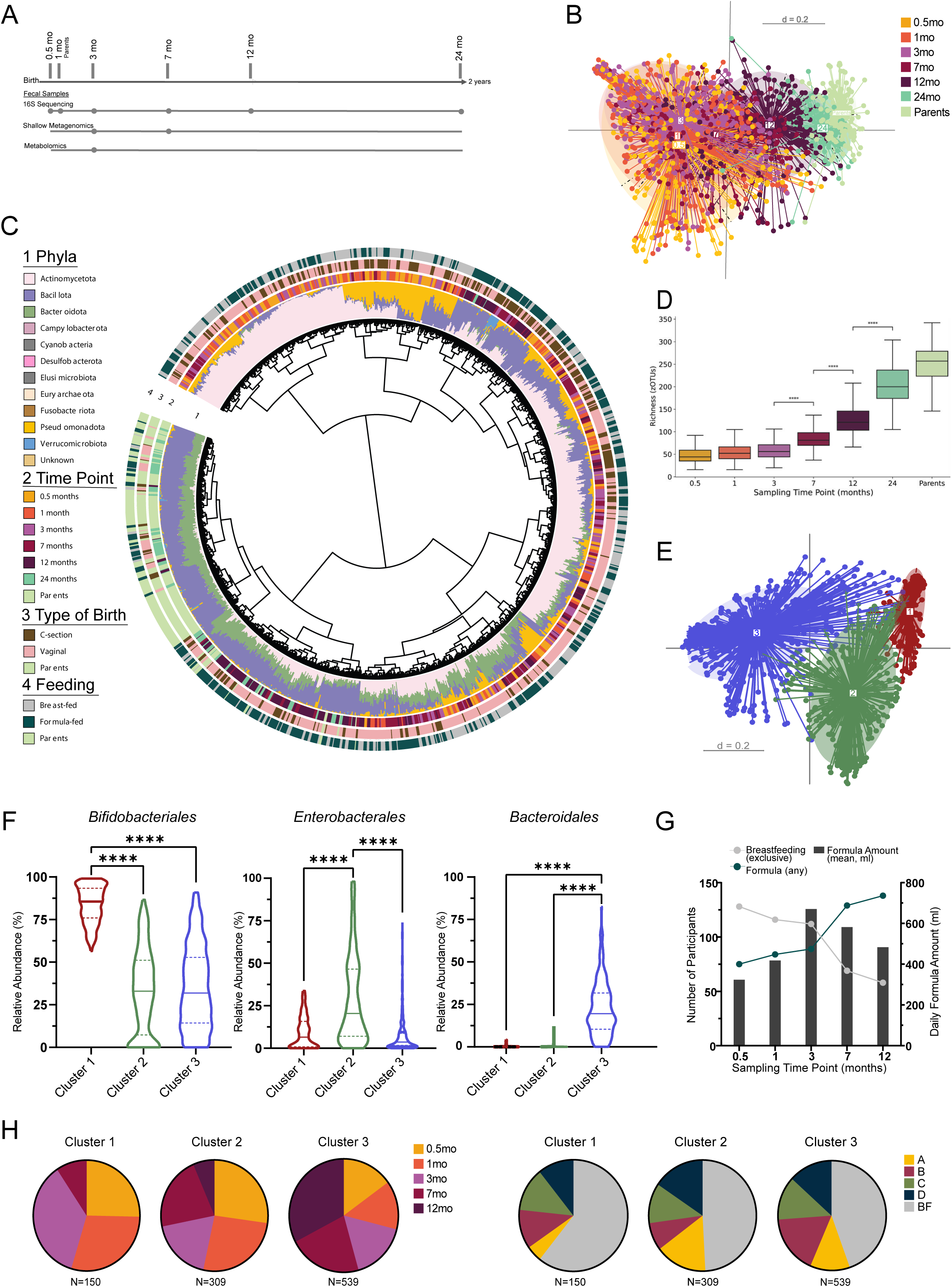
Temporal development of infant microbiota. (A) Schematic of study fecal sample collection and time points of analysis of fecal samples. (B) Non-metric multidimensional scaling (NMDS) plot of fecal samples grouped according to infant age (months) or parent sample (separation of groups as whole is significant: p ≤ 0.001, permutational multivariate analysis based on distance matrices). (C) *De novo* phylogenetic clustering of all study fecal samples (1222 fecal samples from 210 infants (multiple time points, including follow-up) and 152 parents (single time point)) displayed as a dendrogram, phyla per sample as stacked bar plots (second innermost ring), metadata for each sample in the outermost three rings (feed type, type of birth and sampling time point). (D) Alpha-diversity (richness, zOTUs) at different infant sampling time points and for parental samples. (**** = p ≤ 0.0001, 25^th^ to the 75^th^ percentile box plot, median: solid line, whiskers: min to max). (E) Metric multidimensional scaling (MDS) plot of beta-diversity (unsupervised clustering) of all infant samples (n= 998) producing three distinct clusters (separation of groups as whole is significant: p ≤ 0.001, permutational multivariate analysis based on distance matrices). (F) Top 3 relative abundance (%) cluster dominating bacterial order violin plot (**** = p ≤ 0.0001, median: solid line, 25^th^ and 75^th^ percentile: dashed lines), de novo generated clusters. (G) Mean daily formula intake (ml, bar plot) and exclusive breastfeeding rates (grey line) versus formula consumers (turquoise line). (H) Descriptive cluster composition regarding sampling time point and feed type (breastfeeding: grey, formula A (placebo): yellow, formula B (bifidobacteria): red, formula C (GOS): green, formula D (bifidobacteria + GOS): dark blue).

**Table 1.** Cohort characteristics of enrolled infants. Values are given in median with interquartile range in parenthesis unless otherwise indicated.

|  | **A** |  | **B** |  | **C** |  | **D** |  | **BF** |  |
| --- | --- | --- | --- | --- | --- | --- | --- | --- | --- | --- |
| N | 35 | - | 39 | - | 36 | - | 37 | - | 63 | - |
| Samples (n) | 119 | - | 136 | - | 125 | - | 135 | - | 483 | - |
| Discontinue study (N) | 4 | - | 2 | - | 1 | - | 2 | - | 2 | - |
| **Birth characteristics** |  |  |  |  |  |  |  |  |  |  |
| Gestational age (days) | 279.0 | (272.0-284.0) | 277.0 | (271.0-284.0) | 279.5 | (273.5-284.0) | 278.0 | (273.0-282.5) | 278.0 | (270.0-285.0) |
| Weight (g) | 3450 | (3100-3725) | 3250 | (2960-3640) | 3410 | (2903-3893) | 3370 | (3060-3585) | 3360 | (3015-3690) |
| Size (cm) | 52.0 | (50.0-53.0) | 51.0 | (49.0-53.0) | 51.5 | (50.0-53.0) | 51.0 | (50.0-52.5) | 52.0 | (50.0-53.0) |
| Cesaerean delivery, % | 37.1 | - | 25.6 | - | 38.9 | - | 40.5 | - | 27.0 | - |
| Female, % | 51.4 | - | 61.5 | - | 30.6 | - | 58.3 | - | 50.8 | - |
| Male, % | 48.6 | - | 38.5 | - | 69.4 | - | 43.2 | - | 49.2 | - |
| Twin, % | 0 | - | 10.3 | - | 11.1 | - | 10.8 | - | 6.3 | - |
| **Maternal characteristics** |  |  |  |  |  |  |  |  |  |  |
| BMI pre pregnancy, kg/m2, mean | 23.2 |  | 22.7 |  | 22.7 |  | 23.9 |  | 22.0 |  |
| Age at inclusion (years), mean | 34.2 |  | 33.1 |  | 33.3 |  | 32.7 |  | 33.4 |  |
| **Nuclear family characteristics** |  |  |  |  |  |  |  |  |  |  |
| Atopic disease, % | 57.1 | - | 59.0 | - | 50.0 | - | 51.4 | - | 60.3 | - |
| Autoimmune disease, % | 8.6 | - | 2.6 | - | 5.6 | - | 8.1 | - | 11.1 | - |
| **Environment** |  |  |  |  |  |  |  |  |  |  |
| Animal contact, % | 31.4 | - | 38.5 | - | 27.8 | - | 35.1 | - | 30.2 | - |
| Distance to study center (km),mean | 36.60 |  | 36.2 |  | 41.20 |  | 39.9 |  | 41.7 |  |
| Siblings in household, % | 22.9 | - | 30.8 | - | 25 | - | 37.8 | - | 44.4 | - |
| **Sample characteristics** |  |  |  |  |  |  |  |  |  |  |
| Breastfeeding during first year, % (N) | 88.6 | - | 87.2 | - | 91.7 | - | 83.8 | - | 100 | - |
| Exclusive formula-feeding, % (N) | 11.4 | - | 12.8 | - | 8.3 | - | 16.2 | - | 0 | - |
| Daily formula intake (ml) | 480.0 | (200.0-720.0) | 500 | (242.5-700.0) | 400.0 | (220.0-750.0) | 505 | (217.5-657.5) | 0 | 0 |
| Age start solid food (months) | 4.9 | (4.4-5.8) | 4.9 | (4.2-5.4) | 5 | (4.6-5.8) | 4.6 | (4.0-6.1) | 5.5 | (5.0-6.1) |

### Temporal development of infant microbiota

Longitudinal infant and parental fecal sampling (n_i_=998; n_p_=152) demonstrated that the temporal assembly of infant microbiota developed towards the parental microbial composition (**Fig. 2B-D, Suppl. Fig. 2A**). Supervised beta-diversity analysis illustrated significant shifts of microbiota profiles at month 3-24, yet showing clearly distinct patterns compared to parental samples, whereas the early sampling time points (0.5, 1 and 3 months) were characterized by indistinguishable clusters of microbiota profiles (**Fig. 2B**). Unsupervised analysis based on generalized UniFrac distances identified remarkable heterogeneity in microbiota profiles largely independent of the feeding type or birth groups (C-section vs vaginal birth) (**Fig. 2C**). Parental samples appeared in two distinct phylogenetic clusters and were flanked by samples from the older sampling time points with Bacteroidota (synonym Bacteroidetes) and Bacillota (syn. Firmicutes) becoming more dominant. Relative abundance of Actinomycetota (syn. Actinobacteria) was highly variable at early sampling time points and decreased with age (**Fig. 2C**). Gradual development of the infant microbiota is highlighted by increasing richness and alpha-diversity reaching significance at 3 months of age. Significant differences in average individual richness and Shannon effective number of species between infants and parents dissolved after 24 months (**Fig. 2D, Suppl. Fig. 2A**).

Unsupervised hierarchical clustering identified three distinct clusters across all infant samples during the first year of life (**Fig. 2E**). The clusters were characterized by a disproportionally high relative abundance (%) of the orders Bifidobacteriales (Cluster 1), Enterobacterales (Cluster 2) and Bacteroidales (Cluster 3) (**Fig. 2F**). Minor differences in bacterial abundance between the clusters included Eubacteriales (syn. Lachnospirales), Lactobacillales and Veillonellales-Selenomonadales (**Suppl. Fig. 2B**). The Bifidobacteriales*-*dominated cluster was enriched for breast-fed samples (adj. p-value = 0.003) and largely composed of fecal samples from early time points (7 and 12 months underrepresented, adj. p-value = 0.0004 and 3.904e-15, respectively). Formula consumption peaked at 3 months of age and breast-feeding simultaneously decreased with increasing formula milk consumption (**Fig. 2G**). Concordant with decreasing formula intake after 3 months (**Fig. 2G**), 97.5% of infants received solid foods by month 7 (**Table 1**). The two other clusters also included samples from older infants (7 and 12 months) (**Fig. 2H**). Samples from interventions and breast-fed infants (**Fig. 2H**) were equally present in clusters 2 and 3. In addition, fecal samples from babies born via C-section were overrepresented in cluster 2 (adj. p-value = 1.072e-18), and those born vaginally in cluster 3 (adj. p-value = 3.512e-19) (**Suppl. Fig. 2C**).

To define the impact of delivery and feeding modes on transmitted bacterial taxa from mothers to infants, we calculated the ratio of bacterial transmission between the mother-child dyads ordered by type of birth and feeding group. We defined this ratio as the number of shared zOTUs between the mother and the child divided by the total number of zOTUs present in the mother. We observed an increasing trend in the transmissibility over time during the 1^st^ year (Kruskal Wallis p = 2.2e-16, **Suppl. Fig. 2D**). While for each individual time point, the type of feeding (feed groups A to D) did not significantly influence the bacterial transmissibility (data not shown), transmissibility was different between feed groups for all time points combined (p =0.00063). A post-hoc Dunn test showed that BF was significantly different to formula A and B in the 1^st^ year of life (**Suppl. Fig. 2D**). The type of birth significantly affected the bacterial transmission in the 1^st^ year (p = 0.0037). As previously shown^35^, C-section birth altered bacterial transmission from mother at early age (months 0.5 and 1) but this trend was not significant at later time points (months 3, 7 and 12) (**Suppl. Fig. 2E**).

### Impact of bifidobacterial and GOS intervention on infant microbiota assembly

Generalized UniFrac distance showed no significant differences between community structures of the four differentially supplemented infant formulas (A-D) and the exclusively breast-fed group (**Fig. 3A**), suggesting that inter-individual variations largely trumped the stratification of microbiota profiles based on feeding regiments. Species richness increased equally across feeding groups as infants aged (**Fig. 3B**). Beta-diversity was not significantly different between breast-feeders and the formula groups (**Suppl. Fig. 3A),** but differed between the placebo (Formula A) and other supplemented groups at age 3 months (**Suppl. Fig. 3B)**. No differences were seen at 24-months (**Suppl. Fig. 3C**). Beta-diversity was significantly different between babies born vaginally vs. by C-section especially at early age (0.5 and 1 months; **Suppl. Fig. 3D**). Progression in relative abundance of the top ten taxa over time are summarized in the horizon blot (**Fig. 3C**). *Bifidobacterium* was the most abundant genus (41.2%) across all infant fecal samples reaching peak abundance at the age of 3 months (**Fig. 3C**). Average abundance of *Bifidobacterium* was lowest in the placebo group A (32.3%) followed by the bifidobacteria-supplemented group B (39.7%). *Bacteroides, Escherichia-Shigella*, *Veillonella, Klebsiella, Clostridium, Streptococcus* and *Enterococcus* contribute to the very early life infant communities (months 0.5, 1, 3) at substantially lower abundance levels (<13%) (**Fig. 3C**). Relative to the taxon median, the genera *Bacteroides* and *Blautia* clearly increased in average abundance at month 7 and 12 (**Fig. 3C**). Despite some variation over time, formula-fed infants showed on average a 29% higher abundance of *Veillonella* compared to breast-fed infants (**Fig. 3C**). GOS supplementation in the interventional groups C and group D (*Bifidobacterium* and GOS) significantly increased the average bifidobacterial abundance compared to the control group A at month 3 and 7 of age (low formula consumers omitted, **Fig. 3D**) but formula supplementation with *B. longum* and *B. breve* (group B) failed to increase it. The comparison of bifidobacterial abundances between breast-fed and formula fed-infants (low formula consumers omitted) showed a complete loss of significance between any of the interventional groups at 3 months, demonstrating that the inherent variability between samples is higher than the effect of the intervention at this time point. At 7 months, significance of the GOS-mediated effects on increased bifidobacterial abundance was maintained, suggesting that time of exposure to formula supplements is important to sustain effects (data not shown). No additional differences were detected between interventions (data not shown). Shallow metagenomic analysis identified *B. longum*, *B. breve*, *V. parvula*, *E. coli* and *B. bifidum* as the 5 most abundant and prevalent bacterial species at month 3 and 7 independent of the feeding groups (**Fig. 3E**). Some breast-fed infants lacked the most common *Bifidobacterium* strains and instead harbored different *Veillonella* species and *Enterococcus faecalis* (**Fig. 3E**). Correlation analysis based on shallow metagenomics identified pathways related to amino acid biosynthesis (yellow boxes), nucleoside and nucleotide biosynthesis (blue), cell structure biosynthesis (green) and cofactor, carrier and vitamin biosynthesis (orange) in decreasing abundance (**Fig. 3F**). In accordance with the low fiber availability at these early life stages, fermentation was the least abundant pathway. No clear clustering occurred in relation to age and feeding (**Fig. 3F**).

**Figure 3.**
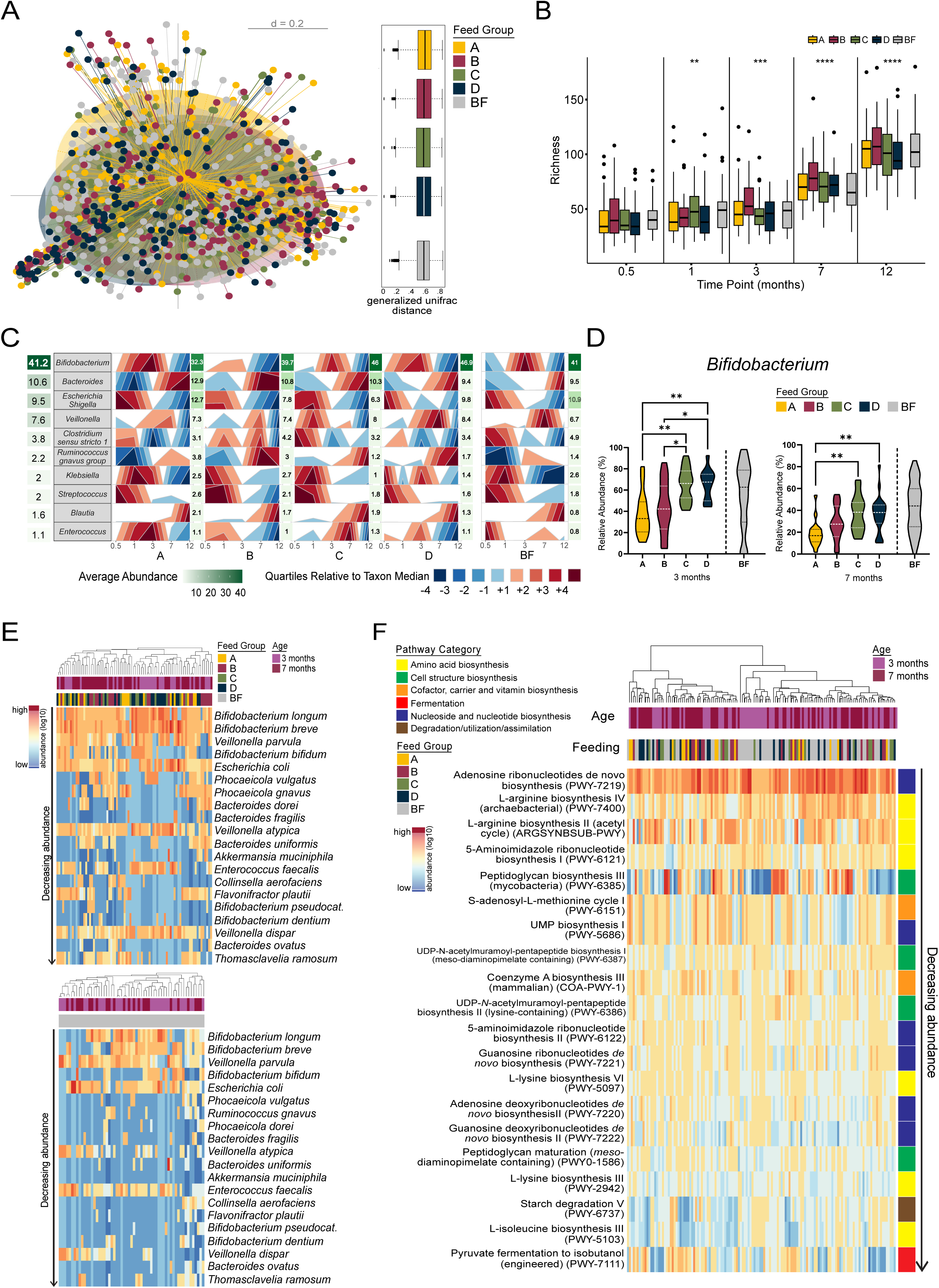
Impact of bifidobacterial and GOS intervention on infant microbiota assembly. (A) Infant beta-diversity (MDS plot) based on feed group. Boxplots show generalized UniFrac distance for each of the five feed groups. (B) Richness over time in the different feed groups (0.5:1 mo: ** = p ≤ 0.01, 0.5:3 mo: *** = p ≤ 0.001, 0.5:7 mo and 0.5:12 mo **** = p ≤ 0.0001, Anova p < 2.2e−16). (C) Top 10 relative abundance (%) taxa (genus) found in >80% of samples (average abundance >= 0.5). Color scale corresponding to increase (red) or decrease (blue) abundance relative to the taxon median, average abundance per feed group in green boxes. (D) Frequency distribution of the relative abundance of the genus *Bifidobacterium* (median: solid line, upper and lower quartile: dotted line, ** = p ≤ 0.01, * = p ≤ 0.05). Minimal formula consumers were omitted (lowest quartile based on daily formula volume consumption). Breast-fed infants are figured on the right for comparison. (E) Relative abundance (log values) of the top 20 species (shallow metagenomics) of individual samples. Metadata as bars above samples (feeding, age) with hierarchal clustering at the very top. (F) Top 20 identified pathways (column normalized) associated with shallow metagenomic sequencing samples sorted by decreasing abundance. Metadata as bars above samples (feeding, age) with hierarchal clustering at the very top; pathway (BioCyc identifier), and pathway categories on the left and right of the heatmap respectively.

### Metabolites clearly differ between feeding groups and display diurnal rhythmicity

We next characterized the metabolite milieu at three months of age since solid food has not been introduced yet and infants consumed the highest volume of formula at this time point. Principal Component Analysis (PCA) identified significantly separated clusters for breast-fed and formula-fed infants, while formula-fed infants remained indistinguishable (**Fig. 4A**, left blot). Interestingly, the exclusively formula-fed infants clustered away from the breast-fed infants, while the mixed feeding (receiving breastmilk and formula) partially overlapped with the breast-fed infants (**Fig. 4A**, right blot), suggesting that the intake of either breastmilk or infant formula plays a significant role in the respective fecal metabolic profiles. To further resolve the metabolome differences between breast-fed and formula-fed infants, we performed additional analyses (PCA) of metabolite profiles from human and formula milk, including 103 breastmilk samples and the 4 different formulas. Here, we demonstrated that formula samples clearly cluster away from all breastmilk samples (**Suppl. Fig. 1D**). Furthermore, breastmilk samples separated in two distinct clusters which equally distribute across all fecal samples (**Suppl. Fig. 1E**), suggesting that the difference in the breastmilk metabolome are not exclusively responsible for the differences in the fecal metabolome.

**Figure 4.**
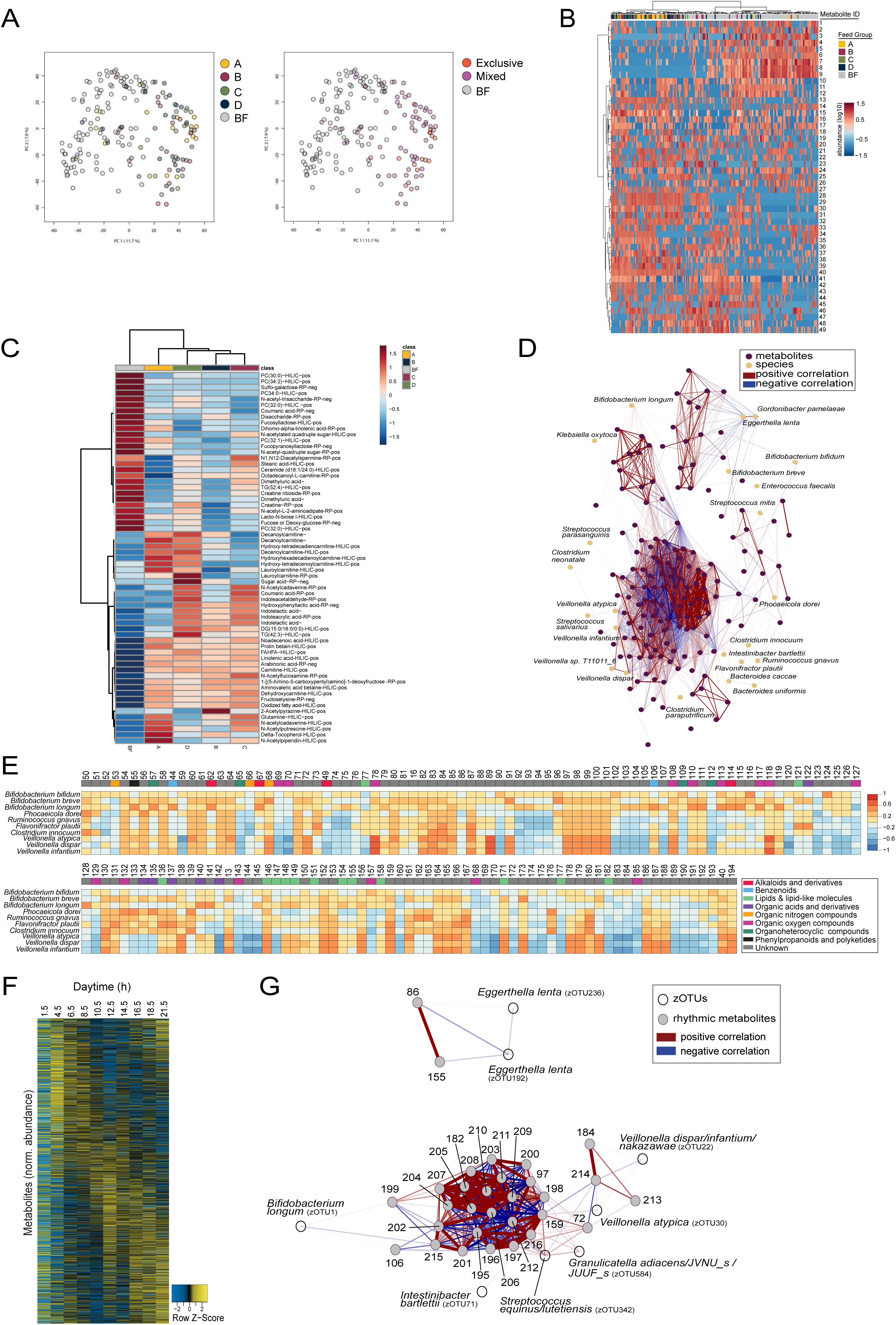
Metabolites clearly differentiate feeding groups and display circadian changes. (A) Principal component analysis of infant metabolite profiles (3 months) stratified according to feed type (left panel) and exclusive *vs* non-exclusive feeders (right panel). (B) Intra-sample-normalized heatmap depicting the relative abundance of the top 49 metabolites found in infant fecal samples (3 months). (C) Mean abundance of annotated fecal metabolites (3 months) displayed per feed group. (D) Correlational network of fecal metabolites and species identified using shallow metagenomics sequencing. (E) Heatmap of significantly correlated (r > 0.5) species (shallow metagenomic sequencing) and fecal metabolites, each metabolite is color-coded according to category affiliation to MetaCyc pathways. (F) Heatmap of normalized abundance of fecal metabolites at time-of-day intervals. Metabolites are sorted by their peak abundance determined with a cosine-wave regression. (G) Network correlational analysis of zOTUs and rhythmic metabolites. Thickness of connecting lines indicates strength of correlation.

Cluster analysis of metabolite abundances was used to stratify patterns according to the feeding groups (**Fig. 4B**). As illustrated in the heatmap for the 49 most abundant features, breast- and formula-fed infants differentially clustered with only occasional appearance of samples from formula-fed infants in the breast-fed cluster (**Fig. 4B**). A more detailed analysis on the respective metabolites, phosphocholines as well as some specialized sugar derivatives like sugar acid, N-acetylated quadruple sugar or N-acetylated trisaccharide are increased in the breast-fed group, whereas carnitines is increased in all formula-fed groups. Indole derivatives such as bacterial-derived indolelactic acid is also increased in formula-fed groups B, C and D (**Fig. 4C**). Network correlational analysis was employed to identify the association of bacterial species and fecal metabolites (**Fig. 4D**). The top ten significantly bacterial species correlated positively or negatively with metabolites, suggesting functional redundancy between taxa (**Fig. 4E**). Variation in abundance of metabolites and bacteria according to the sampling daytime was previously identified in adult populations^27,28^. To evaluate whether this also occurs in infants in the first year of life, we analyzed daytime-dependent fluctuations in metabolite abundance at age 3 months. Peak abundance of fecal metabolites across all fecal samples is illustrated in the heatmap, supporting diurnal rhythms (**Fig. 4F**). Rhythmic metabolites appeared different depending on each feed and birth mode but maintained rhythms (**Suppl. Fig. 4A, B**). Network correlational analysis further correlated bacteria (zOTUs), such as *B. longum* and *Veillonella* and rhythmic metabolites (**Fig. 4G**). These findings support the hypothesis that circadian regulation starts at early stages of life with possible functional contribution to the infant microbiome.

### The development of diurnal rhythmicity of the infant microbiota is affected by age and formula supplementation

In accordance to daytime-dependent differences in metabolite abundance and their link to distinct bacterial taxa (**Fig. 4G**), significant diurnal rhythms were found in bacterial richness and alpha-diversity of fecal samples collected from infants throughout day and night (**Fig. 5A, Suppl. Fig. 4C, D**). Diurnal rhythmicity was also present at the level of individual zOTUs as illustrated by their normalized peak abundance across the 24 h day (**Fig. 5B**). Interestingly, significant rhythmicity was detected for the highly abundant genera *Bifidobacterium* (**Fig. 5C**). The amount of diurnal zOTUs increased as infants mature in parallel to the increasing diversification of infant gut microbiota, with highest numbers of rhythmic zOTUs at age 12 months (**Fig. 5D**). Of note, rhythmic zOTUs were most numerous in cluster 3 containing samples of older infants (**Fig. 2H, Suppl. Fig. 4E**). Rhythmic zOTUs were detected dependent on the birth types (**Suppl. Fig. 4F**). Interestingly, the number of rhythmic bacteria was highest in formula groups supplemented with bifidobacteria and GOS, which even exceeded the number of rhythmic zOTUs in breast-fed infants (**Fig. 5E**). Independent of the groups, the highest numbers of rhythmic zOTUs belonged to the genera *Veillonella* (7 zOTUs), *Bacteroides* (6 zOTUs), *Bifidobacterium* (6 zOTUs), *Streptococcus* (6 zOTUs) and *Clostridium* (5 zOTUs). PICRUSt2 analysis revealed that the total amount of significantly rhythmic assigned pathways dominated in samples from formula group D at age 7 months (604), followed by the breast-fed group showing rhythmicity in 162 pathways (**Fig. 5F**). Furthermore, LEfSe analysis^36^ of these rhythmic metabolic pathways identified that the placebo group (formula A) differed the most from the other feeding groups and particularly 29 rhythmic pathways differed between placebo and breast-fed infants (**Suppl. Fig. 4G**). Interestingly, in all group-to-placebo comparisons, fatty acid beta-degradation was enriched in placebo-fed infants (**Suppl. Fig. 4G**). In contrast to the placebo group, results from the groups supplemented with formula containing bifidobacteria and GOS most closely resembled the breast-fed group.

**Figure 5.**
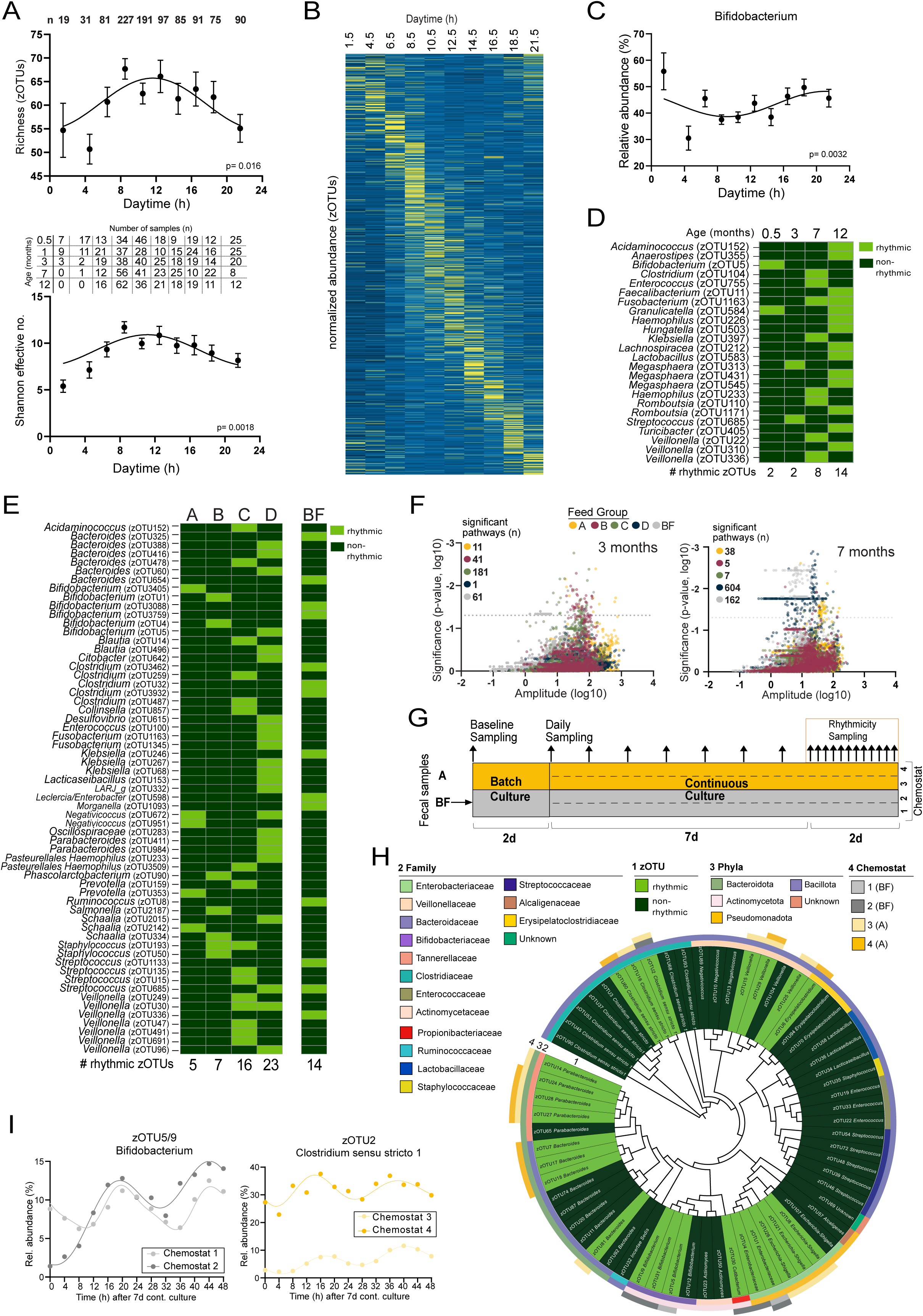
Developing daytime-related rhythmicity of the infant microbiota is affected by formula supplementation. (A) Diurnal profiles of alpha-diversity of samples with known time of defecation (n = 987), displayed as mean ± SEM. Number of total samples at each time point are shown above the graph, number of samples per time of day per age group are displayed in the table in graph. (B) Heatmap of relative abundance of 777 zOTUs detected in infant samples (mean relative abundance > 1%, normalized to peak per zOTU) ordered by time of cosine-wave amplitude peak of the zOTUs. (C) Diurnal profile of the genus *Bifidobacterium* across all infant samples. (D) Rhythmic zOTUs at age 0.5, 3, 7 and 12 months with the total number of rhythmic zOTUs displayed at the bottom of the graph. (E) Rhythmic zOTUs in the five feeding groups with the total number of rhythmic zOTUs displayed at the bottom of the graph. (F) Quantification and significance of rhythmic and arrhythmic pathways in infant samples (3 and 7 months), separated by feed group. Significantly rhythmic pathways are located above the dashed line (p < 0.05). (G) Schematic illustration of study design and sampling time points of chemostats 1-4 containing stool cultures obtained from breastfed (BF) and placebo formula (A) fed infants. (H) Phylogenetic clustering of zOTUs identified in chemostat samples, displayed as a cladogram. Rhythmicity is indicated for the individuals zOTUS (ring 1), family (ring 2), phyla (ring 3) and in which chemostat the rhythmic zOTUs were found (ring 4). (I) Circadian profiles of zOTUs from chemostat cultures after 7 days of continuous culture. Significant rhythms are illustrated with fitted cosine-wave curves (cosine-wave regression, p ≤ 0.05). Data are represented as mean ± SEM.

### Circadian rhythms in dominant infant microbial taxa persist *in vitro*

In order to investigate whether microbial rhythms observed during early life are host-driven or rely on microbial intrinsic clock mechanisms, fecal samples from selected exclusive breast-fed and formula-fed (group A) infants at age 3 months were cultured in a chemostat model. To establish stable growth conditions, an initial adaptation phase (batch culture) was set up followed by continuous culturing for 7 days in the chemostat to ensure host-independent conditions (**Fig. 5G**). Beta diversity was similar between chemostats containing stool samples from the breast-fed donor, but differed significantly between feeding conditions (**Suppl. Fig. 5A**). In contrast to the inoculum, which was highly enriched in *Bifidobacterium* (>90% abundance) in the breast-fed sample, *Bifidobacterium* reached notably lower average abundance in the chemostat (12.79 < 89.73 in breast-fed; 4.74 < 75.49 in group A) (**Suppl. Fig. 5B**). In the second paired chemostat, the inoculum came from placebo formula group A, and the genera *Clostridium* and *Escherichia−Shigella* were similarly enriched, whereas *Bifidobacterium* was low in abundance (**Suppl. Fig. 5B)**. To investigate potential rhythmic alterations in abundance, beginning at day seven of continuous culture, samples were taken every 4 hours over the course of 48 hours (**Fig. 5G)**. Circadian analysis on microbiota composition revealed truly endogenous circadian rhythms in all chemostats illustrated at the level of dominant phyla, families and genera (**Fig 5H, Suppl. Fig. 5B, C**). Rhythmic changes in abundance are highlighted for highly abundant zOTUs (**Fig. 5I, Suppl. Fig. 5D**) and selected metabolic pathways (PICRUSt2 analysis, **Suppl. Fig. 5E**). In accordance with results demonstrated in infant stool samples (**Fig. 5C, E**), significant rhythms were identified for zOTUs of the genera *Bifidobacterium* and *Enterobacteriaceae,* which were overrepresented in the breast-fed samples, and *Clostridium sensu stricto* 1, which were overrepresented in chemostat samples from formula A (**Fig. 5I, Suppl. Fig. 5D**). These results are the first demonstration that specific taxa from a complex microbiota undergo circadian rhythmicity after seven days of continuous culture, pointing to a host-independent, intrinsic establishment of rhythmicity in the infant bacteria.

## Discussion

In this randomized, controlled intervention trial, we demonstrated that formula- and breast-fed infants share characteristic features of microbiota assembly in the first year of life including age-related increase in bacterial richness and diversity as well as dominant abundance of specific bifidobacterial species. Interestingly 24-hour bacterial oscillations were present in all groups and could be integrated into rhythmic metabolite networks. Of note, bacterial profiles highly varied between individuals, even though the study population was homogenous related to geographical area, formula consumption, daily routines (sleep, eat, diaper change) as well as birth, family and health characteristics of the cohort. Interestingly, formula- and breast-fed infants substantially differed in their metabolite profiles, despite considerably overlapping bacterial community structures, emphasizing the unique composition of breast milk.^37^ Nevertheless, and consistent with previous findings,^15,38,39^ GOS supplemented infant formula was most efficient to sustain high levels of the genus *Bifidobacterium* compared to formula containing bifidobacteria or placebo. Remarkably, the presence of GOS in infant formula resulted in the high numbers of rhythmic zOTUs across a range of different bacterial taxa, even exceeding the number of rhythmic zOTUs in breastfed infants. Similar but less pronounced was the effect of formula supplementation with *B. longum* and *B. breve*, despite representing the most abundant species in the infant gut, however placebo showed the fewest number of rhythmic bacterial taxa. Host independent circadian regulation of bacterial rhythmicity was confirmed in breast-fed and infant formula derived stool microbiota using an *ex vivo* gut chemostat model. These findings demonstrate for the first time that bacterial intrinsic clock mechanisms contribute to the development of circadian rhythmicity in the microbiota at early life stages.

The infant microbiome gradually develops towards a more diverse adult-like microbiome over time.^7^ Interestingly, shallow metagenomics sequencing of samples from 3- and 7-months old infants demonstrated that *Bifidobacterium* (*B. breve*, *B. longum* and *B. bifidum*), *Veillonella* and *Enterobacteriaceae* dominate this early life stage, irrespective of the diet received. These results are consistent with previously published baby cohort where fecal samples of 40 days, 3-months and 6-months old infants were compared, showing that *Bifidobacterium* and *Enterobacteriaceae* were the most predominant taxa, regardless of diet and age.^40^ Alpha-diversity significant increased from age 3 months onwards, gradually approaching parental diversity levels at age 24 months. This is in line with previous reports indicating significant changes in the dominant phyla and diversity appearing from age 3 months onwards^16^ or at the end of the first year of life^41^ until the parent’s diversity and composition is reached by month 14^16^ or on the second birthday.^42^ None of the three types of supplementation investigated in this study were able to fully recreate the breast milk-related microbial environment. Accordingly, the fecal metabolic profile of 3-months old, breast-fed babies was clearly distinguishable from formula-consuming infants. Untargeted metabolomics of breastmilk and infant samples identified clear differences in the metabolite profiles and considering the respective fecal samples, we conclude that the intake of either breastmilk or infant formula plays a significant role in shaping the fecal metabolome. Breast-fed babies showed a high number of organic oxygen metabolites in their feces, whereas formula-fed babies were associated with a lipid-enriched metabolome composition. Similarly, metabolic profiles of 12 month old infants were reported to be distinguishable between diets.^43^ In addition, the two distinct clusters of metabolite profiles in breastmilk, which appeared at least in part to be differentiated by the presence of the human milk oligosaccharide fucosyllactose, are not exclusively responsible for the differences in the fecal metabolome.

Previous work in adult populations, done by others as well as our group, documented that microbiota composition and function undergoes diurnal rhythmicity.^26,44^ However, whether these rhythms appear at early age remained unknown. We provide the first evidence that during the first year of life in all feeding groups, the abundance of the most dominant taxa, including *Bifidobacterium*, varied depending on the time of defecation and circadian rhythm analysis revealed that these fluctuations follow the 24-hour day. These diurnal oscillations could indeed explain the opposing results on abundance fluctuations reported, e.g. for bifidobacteria^45^ if samples were taken at different day times in previous studies. Here we discovered significant daytime-dependent fluctuations in bacterial diversity, at the level of genera and zOTUs in infant stool samples. Although rhythmicity was present in all feeding groups and at all ages examined, including the genera *Veillonella*, *Bacteroides*, *Bifidobacterium*, *Streptococcus* and *Clostridium*, dramatic differences were detected between rhythmic zOTUs based on infant, age and diet. The overall highest number of rhythmic zOTUs was found in infants fed with formula supplemented with the combination of bifidobacteria and GOS, which exceeded numbers seen in breast-fed infants. Moreover, circadian analysis identified the presence of diurnal rhythms in microbiota composition and function already in 2-weeks old infants demonstrating that microbial oscillations are an early event in the infant gut development. The number of rhythmic bacteria gradually increased as the infant gut matured and regular diet was introduced to the infant nutrition.

Circadian regulation of microbiota composition and function was recently shown to depend on the host circadian clock, particularly in intestinal epithelial cells.^46^ Conversely, microbiota have been shown to influence host circadian rhythms,^47,48^ indicating a cross-talk between the host interface and bacteria. Microbiota transfer experiments further demonstrated the physiological relevance of microbial oscillations for the host’s metabolic and gastrointestinal health.^26,46,49^ Since environmental factors, such as the light-dark-cycle, or time of food intake, are potent signals impacting microbial rhythms,^26,46,49,50^ abnormal light or food exposure can alter microbial oscillations and thus might lead to the development of pathologies and disease. To gain further insights into the possible functionality of microbiota oscillations in infants, rhythmic zOTU-associated pathways were examined. Similar to results obtained from the taxonomic analysis, most rhythmic assigned pathways were found in 3- and 7-months old infants fed with formula containing GOS or both the prebiotic supplement and bifidobacteria respectively, followed by breast-fed infants. Importantly, results from the placebo group differed the most from breast-fed infants, whereas results from infants fed with formula containing bifidobacteria and GOS most closely resembled the breast-fed group. The association between the bifidogenic effect of GOS supplementation and the augmentation of the number of rhythmic pathways needs to be further investigated in future studies.

Recent work on single bacterial cultures discovered rhythmic gene expression in *Bacillus subtilis*^24^ and recorded circadian rhythms in the swarming behavior of the gut bacterium *Klebsiella aerogenes*^25^. To identify whether the diurnal rhythms observed in infants gut microbiota are generated by the host circadian clock or bacterial intrinsic mechanism, we cultured complex microbiota communities in a chemostat model. For this purpose, samples from either exclusively breastfed or formula A-fed infants were cultured and supplemented with media containing infant formula. Of note, the relative abundance of taxa as well as bacterial diversity changed significantly when compared to the original inoculum. Despite both inocula being dominated by bifidobacteria, their relative abundance decreased after the continuous flow with culture media begun until a fluctuating but steady abundance was achieved. Importantly, after 7 days *in vitro* when the communities were established, circadian rhythm analysis of microbiota composition provides the first evidence of circadian rhythms in the abundance of dominant bacteria, such as *Bifidobacterium*, *Veillonella*, *Clostridium* and *Bacteroides*, in complex human microbiota communities. Since oscillations from 3-months old infant stool samples are documented outside and independent of the host, these are truly endogenous circadian rhythms and therefore must be generated by bacterial intrinsic clock mechanisms. Importantly, we observed that rhythmic bacterial taxa differed depending on whether the original sample was obtained from an exclusively breastfed infant or exclusively fed with formula A. For example, the composition of breast-fed infants was dominated by rhythmic oscillations of zOTUs belonging to *Enterobacteriacea* and *Bifidobacterium*, whereas formula A-fed samples were enriched with rhythmic zOTUs belonging to *Clostridium* and *Bacteroides.* However, independent of which taxa dominated in both extreme infant feeding exposures (exclusive BF and exclusive formula A), all of them had the potential to become rhythmic *in vitro*, suggesting the importance of bacterial intrinsic clock mechanism for the establishment of circadian microbial communities in the developing infant gut.

Possible limitations of the circadian analysis include the fact that the number of fecal samples collected during the night is lower compared to daytimes and decreases with age. This possibly results from the reluctance of parents to sample feces at night, but also shows the adaptation of older infants to the rhythms of their parents. In any case, the statistical model reached significance even with reduced sample numbers emphasizing the robust finding that distinct taxa of the infant microbiota develop circadian rhythms at early life stages. Furthermore, the explicit contribution of breastmilk relative to infant formula on bacterial rhythms remains unclear, since breastfeeding being the official recommended diet for infants was advocated, and consequently, the group size for exclusively formula-fed infants was limited. Finally, the *ex-vivo* chemostat model allowed us to replicate the infant colonic environment to some extent and to identify bacterial rhythms independent of the host. However, the composition of the infant microbiota was not fully recovered in the chemostat model, suggesting that we only identified an incomplete repertoire of rhythmic taxa. Mechanisms explaining endogenous triggers for bacterial rhythms, such as circadian clock genes, are still missing and will be further investigated.

In summary, this controlled intervention study provided compelling evidence of circadian rhythmicity in the developing infant gut microbiota affected by diet. Our findings warrant the need for further analysis of circadian fluctuations of both bacteria and metabolites and their functional role in contributing to the benefits of infant nutrition.

## Additional information

## Acknowledgements

The Infantibio-II study was initiated and financed by Töpfer GmbH (Dietmannsried, Germany). Besides manufacturing the study formula, they had no role in the conduct and management of the study, analysis and interpretation of data nor the creation of this manuscript. D.H. and S.K. received funding from the German Research Foundation (DFG, Deutsche Forschungsgemeinschaft) SFB 1371 (no. 395357507). D.H. also received funding in the frame of the Joint Programming Initiative of the European Union (project name EcoBiotic) and the German Ministry of Education and Research (BMBF; FKZ 01EA2207). The Technical University of Munich provided technical support through the Core Facility Microbiome of the ZIEL Institute for Food & Health (Klaus Neuhaus, Lukas Mix, Angela Sachsenhauser and Caroline Ziegler) for 16S rRNA gene amplicon sequencing. We thank Nikolai Köhler for giving us advice on integrative data analysis as well as Katharina Sontheimer, Claudia Seegerer and Franziska Kummert for support with sample management and handling. We are most grateful to the participating families and infants of the Infantibio-II study for their time, effort and enthusiasm in our study.

## Author contributions

D.H. conceived and coordinated the project and secured funding. S.K. performed and supervised the circadian analysis. N.H. and D.H. were responsible for creation of trial documents and ethics approval, N.H. was responsible for participant recruitment, enrolment, data collection, management and coordination of collection of all trial samples. N.H., S.R., H.O., S.K. and M.VM. performed 16S rRNA sequencing and subsequent analysis with support from L.S. and M.L., and M.G., S.R. C.M. and K.K. were responsible for metabolomics analysis. S.R. and M.S. performed shotgun metagenomics analysis, H.O., M.H. and M.VM. performed the chemostat experiments and analysis, N.H., S.R., and M.L. performed bioinformatics analysis. D.H., S.K. and H.O. supervised the work and data analysis. N.H., S.K. and D.H. wrote the manuscript. All authors reviewed the manuscript.

---

**Supplemental Figure 1:**
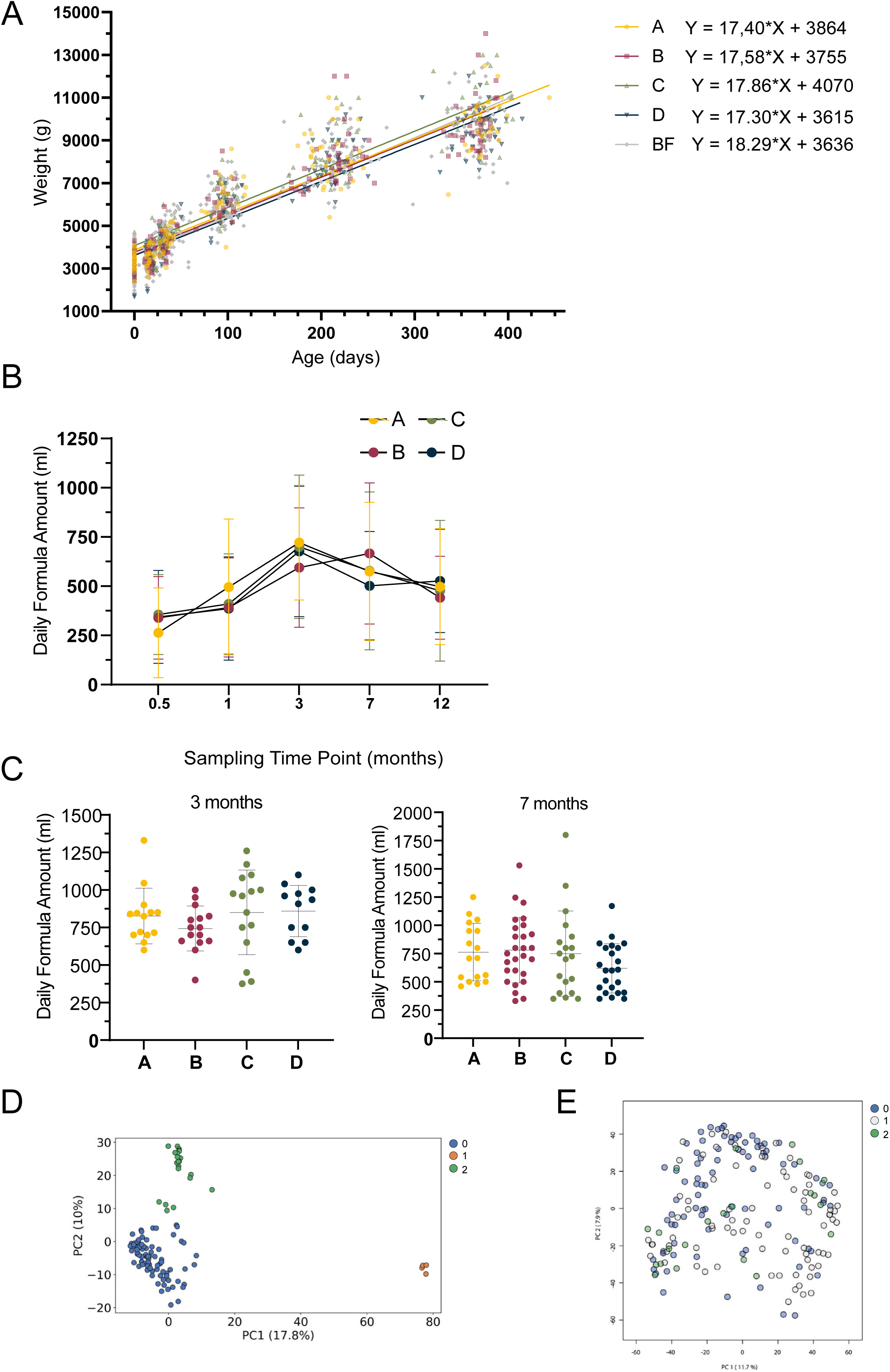
Intervention groups and breastfeeders are comparable with regards to weight gain and formula consumption. (A) Longitudinal weight gain of study participants in the different feeding groups (linear regression, slopes are not significantly different). (B) Daily amount of formula (ml) consumed (mean with SD) per interventional group at different sampling time points. (C) Formula amount consumption (ml) of infants where minimal formula consumers were omitted (lowest quartile based on daily formula volume consumption) at age 3 and 7 months (mean, SD). (D) Principal component analysis of 103 breastmilk samples (1-month), with unsupervised clustering (k-means) by color. t-Test (clusters 0 and 2): 306 significant metabolites on 1439 total (p-value threshold 0.05). (E) Principal component analysis of infant metabolite profiles (3-months) in feces, with breastmilk clusters differentiated by color. t-Test showed no significance between clusters among 4895 metabolites (p-value threshold 0.05).

**Supplemental Figure 2:**
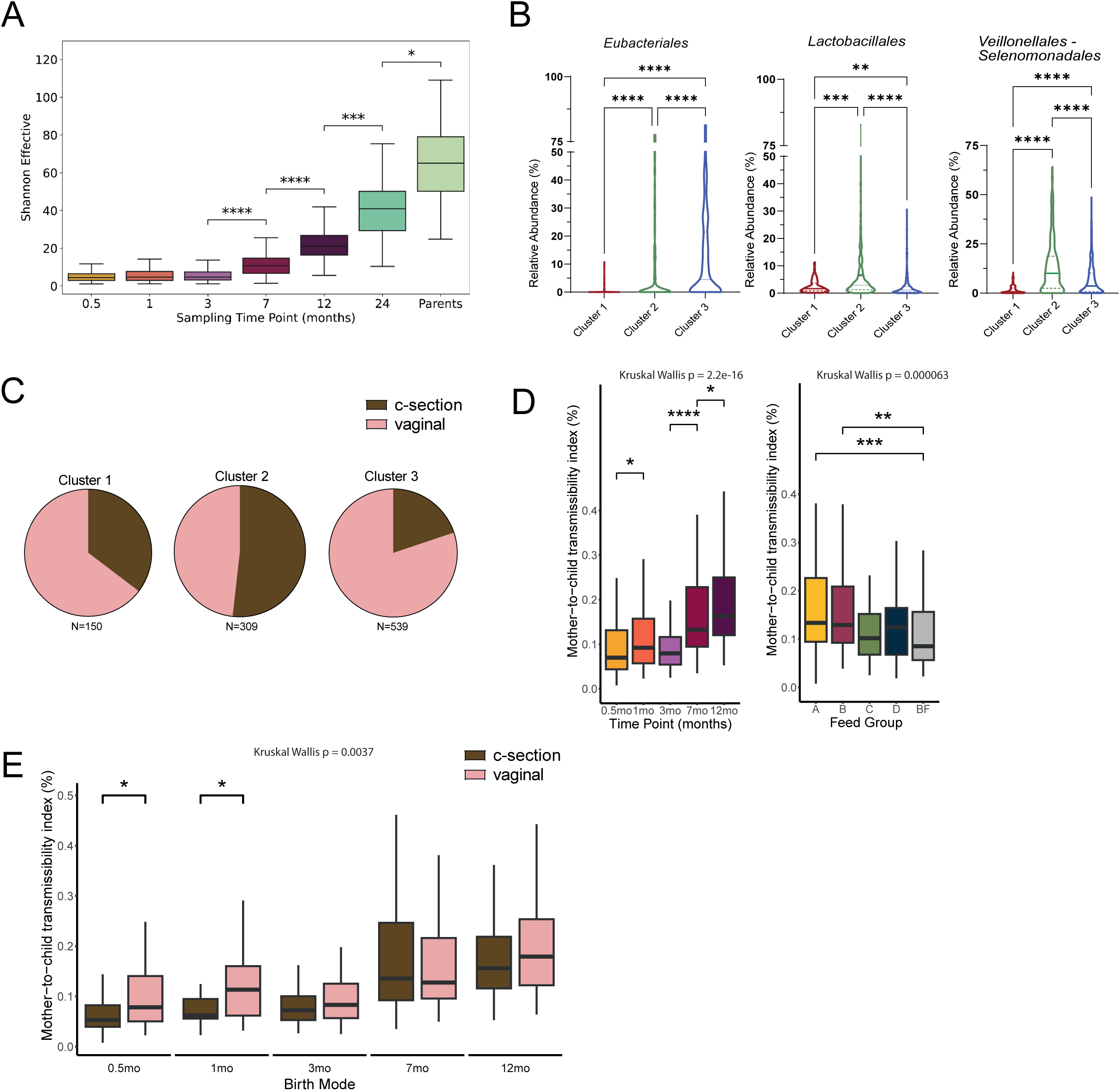
Changes of infant fecal microbiota in relation to age, birth mode and feeding type. (A) Alpha-diversity (Shannon effective number) at all infant sampling time points (including 24 months) and for parental samples (**** = p ≤ 0.0001, 25th to the 75th percentile box plot, median: solid line, whiskers: min to max). (B) Top 4-6 relative abundance (%) cluster dominating bacterial order violin plot (**** = p ≤ 0.0001, median: solid line, 25^th^ and 75^th^ percentile: dashed lines), de novo generated clusters. (C) Birth mode (C-section vs. vaginal) distribution in the three clusters. (D, E) Bacterial (zOTUs) transmissibility ratio (%) between mother and infant over time during the 1^st^ year of life (D, left panel) and according to the feed group (D, right panel) or according to sampling time and birth mode (E). (Dunn test: * = p<0.05, ** = p<0.01, *** = p<0.001, **** = p<0.0001)).

**Supplemental Figure 3:**
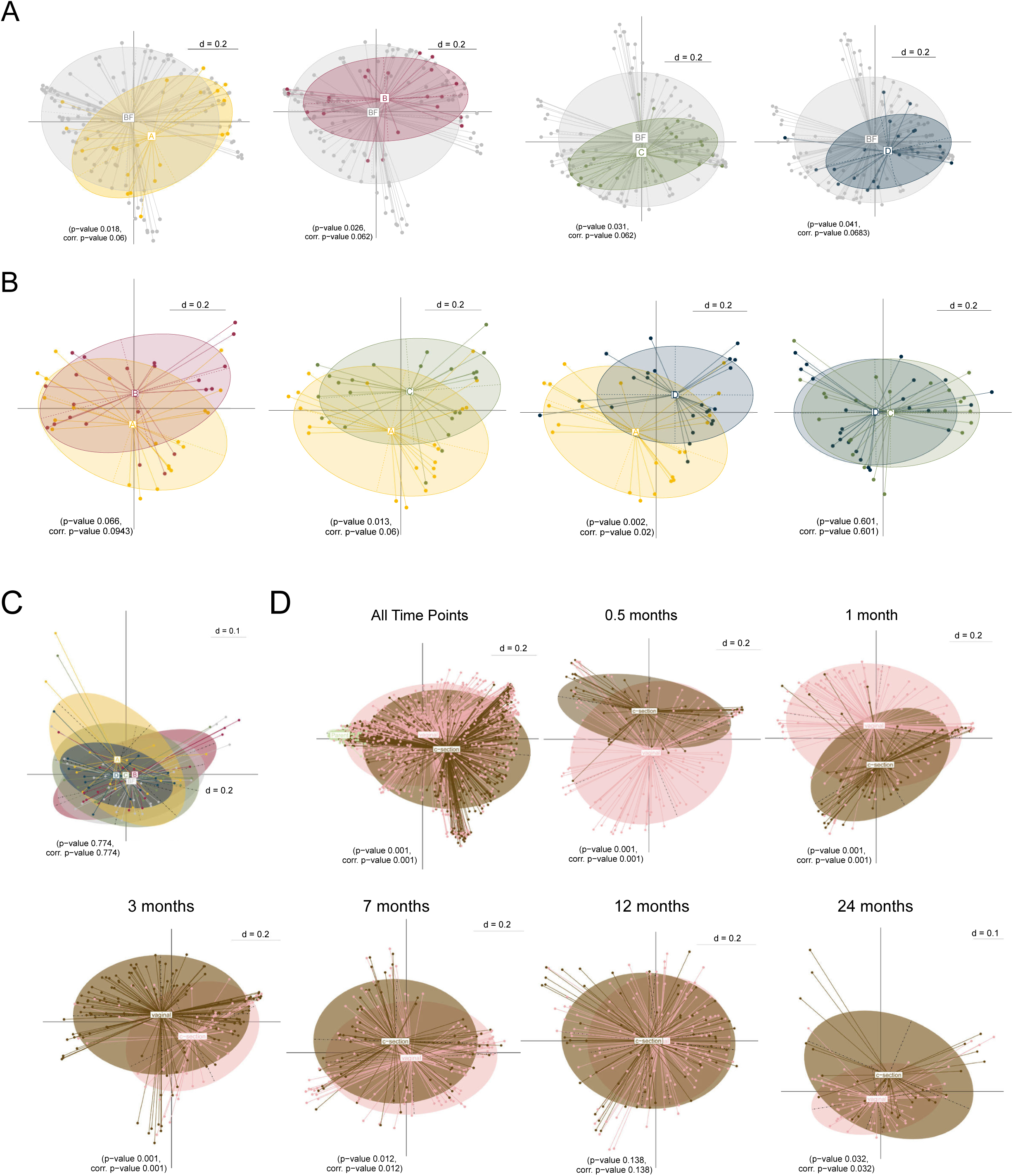
Bacterial profiling in response to feeding and birth mode. (A) MDS plots of beta-diversity comparison between breastfeeders and formula groups at 3 months. (B) MDS plots of beta-diversity comparison between placebo formula A and other groups at 3 months and between C and D. (C) MDS plot of beta-diversity comparison between all feed groups at 24-months. (D) MDS plots of beta-diversity comparisons between vaginal and C-section birth modes overall in the 1^st^ year of life and at each time point.

**Supplemental Figure 4:**
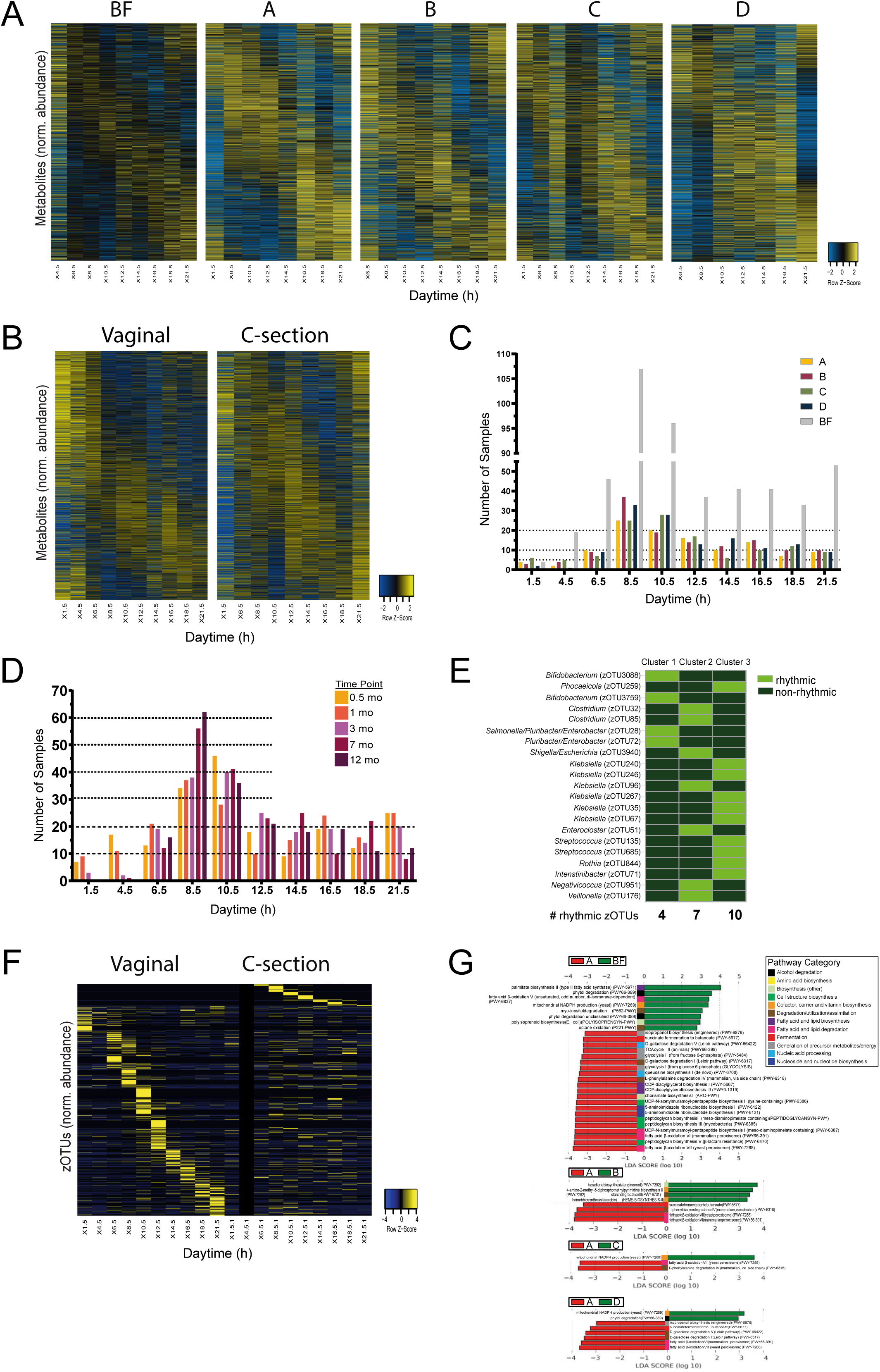
Differences in daytime sampling and rhythmicity in unsupervised clusters and between feed groups and birth mode. (A, B) Heatmaps of normalized abundance of fecal metabolites at time-of-day intervals for each feeding group (A) (1822 metabolites) and birth mode (B) (2317 metabolites). Metabolites were sorted by their peak abundance determined with a cosine-wave regression in each of the groups. (C) Number of fecal samples at different times of day (hours) per feed group. (D) Number of fecal samples at different times of day (hours) per sampling time point. (E) Rhythmic zOTUs in the three de novo clusters with the total number of rhythmic zOTUs displayed at the bottom of the graph. (F) Heatmap of normalized abundance of 948 zOTUs at time-of-day intervals by birth mode. zOTUs were sorted by their peak abundance determined with a cosine-wave regression in the C-section and then the vaginal group. (G) Linear discriminant analysis (LDA) effect size (LEfSe) of pathways at age 7 months between placebo formula group (A) and all other feeding groups (formula B, C, D and breastfeeding (BF)).

**Supplemental Figure 5:**
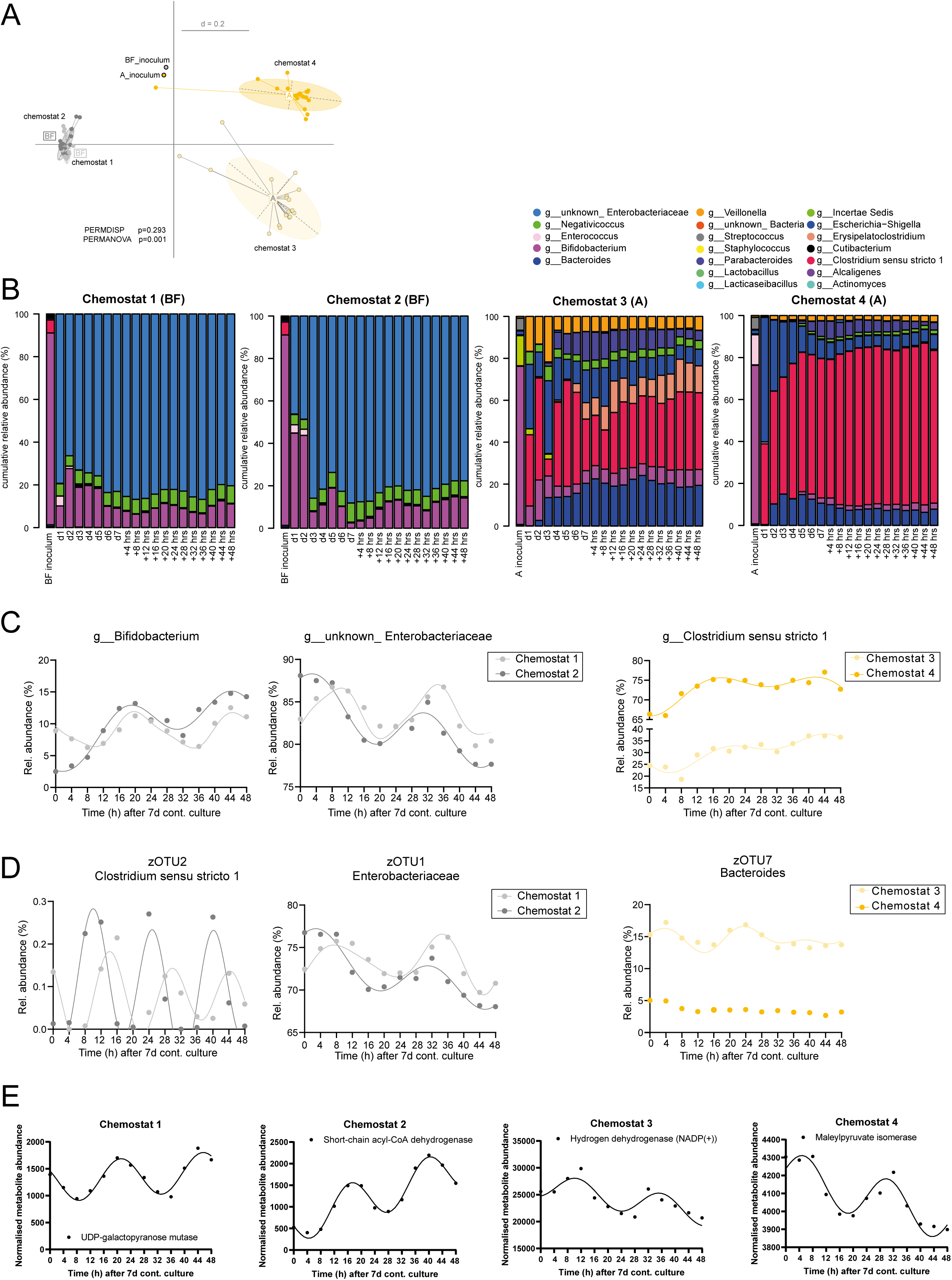
Chemostats differ in taxonomic composition and diversity but consistently show rhythmicity in all vessels. (A) MDS plot of beta-diversity comparison between placebo formula A and BF of the *ex vivo* chemostat samples and the inoculum samples (separation of groups as whole is significant: p ≤ 0.001, permutational multivariate analysis based on distance matrices). BF_chemostat_1-BF_chemostat_2: corr. p-value 0.0627, BF_chemostat_1-A_chemostat_3: corr. p-value 0.003, BF_chemostat_1-A_chemostat_4: corr. p-value 0.003, BF_chemostat_2-A_chemostat_3: corr. p-value 0.003, BF_chemostat_2-A_chemostat_4: corr. p-value 0.003, A_chemostat_3-A_chemostat_4: corr. p-value 0.003 (B) Taxonomic binning barplots showing the differences in taxonomic composition of each chemostat throughout the continuous culture and rhythmicity sampling period. (C-E) Circadian profiles of genera (C), zOTUs (D) and metabolic pathways (PICRUSt2) (E) from chemostat cultures after 7 days of continuous culture. Significant rhythms are illustrated with fitted cosine-wave curves (cosine-wave regression, p ≤ 0.05). For zOTU7, chemostat 4 is not significantly rhythmic. Data are represented as mean ± SEM.

**Supplemental Table 1:** untargeted metabolomics.

| Nr in Figure | Fig | Original ID | RT [sec] | m/z | Compound Name |
| --- | --- | --- | --- | --- | --- |
| 1 | 4C/4D | ID1878_RP_pos | 445.2965 | 584.2579 | unknown |
| 2 | 4C/4D | ID2147_HILIC_pos | 41.3172 | 872.7654 | TG(52:4) |
| 3 | 4C/4D | ID823_RP_neg | 47.9137 | 855.3091 | unknown |
| 4 | 4C/4D | ID1240_RP_pos | 45.7357 | 381.0789 | Disaccharide |
| 5 | 4C/4D | ID1843_HILIC_pos | 46.7741 | 732.5525 | PC(32:1) |
| 6 | 4C/4D | ID1295_HILIC_pos | 325.4208 | 527.3134 | unknown |
| 7 | 4C/4D | ID1979_RP_pos | 47.5239 | 708.2544 | N-acetyl-quadruple sugar |
| 8 | 4C/4D | ID2155_HILIC_pos | 530.2239 | 878.2997 | unknown |
| 9 | 4C/4D | ID2205_HILIC_pos | 569.4872 | 1387.4841 | unknown |
| 10 | 4C/4D | ID666_HILIC_pos | 265.5851 | 337.2509 | unknown |
| 11 | 4C/4D | ID666_RP_neg | 520.0732 | 539.2167 | unknown |
| 12 | 4C/4D | ID784_RP_neg | 519.3508 | 805.5629 | unknown |
| 13 | 4C/4D/4F | ID338_HILIC_pos | 45.5292 | 213.1824 | unknown |
| 14 | 4C/4D | ID947_RP_pos | 645.0067 | 325.2108 | unknown |
| 15 | 4C/4D | ID554_HILIC_pos | 41.3418 | 297.2767 | Noadecenoic acid |
| 16 | 4C/4D/4F | ID1067_RP_pos | 449.214 | 345.2824 | unknown |
| 17 | 4C/4D | ID194_RP_neg | 632.1141 | 295.189 | unknown |
| 18 | 4C/4D | ID1495_HILIC_pos | 466.705 | 604.2497 | unknown |
| 19 | 4C/4D | ID1524_RP_pos | 671.0764 | 430.3485 | unknown |
| 20 | 4C/4D | ID1462_HILIC_pos | 345.6149 | 591.3413 | unknown |
| 21 | 4C/4D | ID203_HILIC_pos | 44.6892 | 165.0937 | unknown |
| 22 | 4C/4D | ID951_HILIC_pos | 43.9798 | 424.3549 | unknown |
| 23 | 4C/4D | ID1501_HILIC_pos | 276.4828 | 604.5999 | unknown |
| 24 | 4C/4D | ID1075_RP_pos | 361.9261 | 351.1558 | unknown |
| 25 | 4C/4D | ID1989_RP_pos | 708.0827 | 763.6034 | unknown |
| 26 | 4C/4D | ID276_RP_neg | 544.2773 | 375.33 | unknown |
| 27 | 4C/4D | ID1872_HILIC_pos | 45.5417 | 747.5897 | unknown |
| 28 | 4C/4D | ID279_RP_pos | 587.4899 | 182.161 | unknown |
| 29 | 4C/4D | ID528_RP_pos | 587.51 | 249.2476 | unknown |
| 30 | 4C/4D | ID1448_HILIC_pos | 44.9654 | 583.5245 | DG(15:0/18:0/0:0) |
| 31 | 4C/4D | ID932_RP_pos | 560.2321 | 322.2425 | unknown |
| 32 | 4C/4D | ID328_HILIC_pos | 45.7918 | 209.187 | unknown |
| 33 | 4C/4D | ID425_HILIC_pos | 70.027 | 255.0003 | unknown |
| 34 | 4C/4D | ID52_RP_pos | 882.4914 | 97.9682 | unknown |
| 35 | 4C/4D | ID1543_RP_pos | 298.375 | 433.2828 | unknown |
| 36 | 4C/4D | ID579_RP_neg | 36.9573 | 502.914 | unknown |
| 37 | 4C/4D | ID957_HILIC_pos | 44.3644 | 425.3386 | Delta-Tocopherol |
| 38 | 4C/4D | ID131_HILIC_pos | 409.5396 | 144.0998 | Prolin betain |
| 39 | 4C/4D | ID962_HILIC_pos | 78.185 | 426.1919 | unknown |
| 40 | 4C/4D/4F | ID2104_HILIC_pos | 44.2302 | 832.669 | unknown |
| 41 | 4C/4D | ID1884_HILIC_pos | 107.927 | 755.6154 | TG(42:3) |
| 42 | 4C/4D | ID1586_HILIC_pos | 130.8504 | 635.6108 | unknown |
| 43 | 4C/4D | ID2017_HILIC_pos | 129.2633 | 801.6593 | unknown |
| 44 | 4C/4D/4F | ID205_RP_pos | 276.8442 | 165.0539 | Coumaric acid |
| 45 | 4C/4D | ID668_RP_pos | 254.763 | 280.1055 | unknown |
| 46 | 4C/4D | ID1138_HILIC_pos | 396.2653 | 472.1415 | unknown |
| 47 | 4C/4D | ID519_RP_neg | 439.0721 | 474.2547 | unknown |
| 48 | 4C/4D | ID136_RP_pos | 68.6502 | 145.1328 | N-Acetylcadaverine |
| 49 | 4C/4D/4F | ID719_RP_pos | 47.173 | 287.241 | N1,N12-Diacetylspermine |
| 50 | 4F | ID13_RP_pos | 45.6086 | 58.0649 | unknown |
| 51 | 4F | ID32_RP_pos | 48.5084 | 72.0808 | unknown |
| 52 | 4F | ID61_RP_pos | 43.7121 | 104.1066 | unknown |
| 53 | 4F | ID98_RP_pos | 47.0312 | 132.0761 | Creatine |
| 54 | 4F | ID125_RP_pos | 587.6851 | 143.1058 | unknown |
| 55 | 4F | ID143_RP_pos | 276.9252 | 147.0429 | Coumarin (low hit rate 48.72%) |
| 56 | 4F | ID170_RP_pos | 587.6044 | 155.178 | unknown |
| 57 | 4F | ID184_RP_pos | 337.5045 | 160.0748 | Indoleacetaldehyde |
| 58 | 4F | ID200_RP_pos | 587.5189 | 164.1504 | unknown |
| 59 | 4F | ID250_RP_pos | 507.8812 | 174.1303 | unknown |
| 60 | 4F | ID262_RP_pos | 587.4744 | 178.1659 | unknown |
| 61 | 4F | ID300_RP_pos | 587.5718 | 186.1558 | unknown |
| 62 | 4F | ID305_RP_pos | 337.5045 | 188.07 | Indoleacrylic acid |
| 63 | 4F | ID318_RP_pos | 587.5566 | 192.1818 | unknown |
| 64 | 4F | ID336_RP_pos | 587.5244 | 196.1768 | unknown |
| 65 | 4F | ID340_RP_pos | 268.5427 | 197.0664 | Dimethyluric acid |
| 66 | 4F | ID365_RP_pos | 47.7627 | 204.0862 | N-acetyl-L-2-aminoadipate |
| 67 | 4F | ID375_RP_pos | 338.6819 | 206.0805 | Indolelactic acid |
| 68 | 4F | ID377_RP_pos | 43.3248 | 206.1021 | N-Acetylfucosamine |
| 69 | 4F | ID515_RP_pos | 47.3348 | 247.0927 | unknown |
| 70 | 4F | ID586_RP_pos | 47.2125 | 264.1182 | Creatine riboside |
| 71 | 4F | ID591_RP_pos | 591.3177 | 265.2523 | unknown |
| 72 | 4F/4H | ID637_RP_pos | 43.2368 | 273.1433 | unknown |
| 73 | 4F | ID699_RP_pos | 38.3207 | 284.9259 | unknown |
| 74 | 4F | ID782_RP_pos | 709.7177 | 299.1386 | unknown |
| 75 | 4F | ID797_RP_pos | 709.6879 | 300.1412 | unknown |
| 76 | 4F | ID814_RP_pos | 374.105 | 302.2304 | unknown |
| 77 | 4F | ID843_RP_pos | 655.1938 | 307.2623 | Dihomo-alpha-linolenic acid |
| 78 | 4F | ID853_RP_pos | 43.1228 | 309.1652 | 1-[(5-Amino-5-carboxypentyl)amino]<br>-1-deoxyfructose |
| 79 | 4F | ID899_RP_pos | 408.5222 | 316.2476 | Decanoylcarnitine |
| 80 | 4F | ID918_RP_pos | 587.8349 | 318.2994 | unknown |
| 81 | 4F | ID1062_RP_pos | 449.149 | 344.2793 | Lauroylcarnitine |
| 82 | 4B/4F | ID1148_RP_pos | 269.0495 | 367.1495 | unknown |
| 83 | 4B/4F | ID1182_RP_pos | 279.9274 | 371.2175 | unknown |
| 84 | 4F | ID1195_RP_pos | 279.7921 | 372.2206 | unknown |
| 85 | 4F | ID1249_RP_pos | 47.249 | 383.1374 | unknown |
| 86 | 4F/4H | ID1276_RP_pos | 424.3017 | 386.2411 | unknown |
| 87 | 4F | ID1299_RP_pos | 279.4737 | 389.2283 | unknown |
| 88 | 4F | ID1385_RP_pos | 385.0854 | 402.283 | unknown |
| 89 | 4F | ID1706_RP_pos | 44.7933 | 471.2175 | unknown |
| 90 | 4F | ID1843_RP_pos | 437.6435 | 552.3348 | unknown |
| 91 | 4F | ID1858_RP_pos | 275.0814 | 569.2906 | unknown |
| 92 | 4F | ID1867_RP_pos | 445.3817 | 583.2545 | unknown |
| 93 | 4F | ID1882_RP_pos | 709.7493 | 585.2672 | unknown |
| 94 | 4F | ID1907_RP_pos | 709.232 | 604.2229 | unknown |
| 95 | 4F | ID1914_RP_pos | 709.8671 | 607.2511 | unknown |
| 96 | 4F | ID1923_RP_pos | 548.5101 | 617.3865 | unknown |
| 97 | 4F/4H | ID1932_RP_pos | 587.3156 | 639.485 | unknown |
| 98 | 4F | ID1948_RP_pos | 587.6053 | 657.459 | unknown |
| 99 | 4F | ID1953_RP_pos | 587.5339 | 658.462 | unknown |
| 100 | 4F | ID1955_RP_pos | 587.5718 | 659.4563 | unknown |
| 101 | 4F | ID1959_RP_pos | 587.5208 | 660.4575 | unknown |
| 102 | 4F | ID4_RP_neg | 44.3665 | 88.0397 | unknown |
| 103 | 4F | ID54_RP_neg | 45.4521 | 162.0766 | unknown |
| 104 | 4F | ID56_RP_neg | 49.2802 | 163.0616 | Fucose or Deoxy-glucose |
| 105 | 4F | ID57_RP_neg | 71.257 | 163.0615 | unknown |
| 106 | 4F/4H | ID58_RP_neg | 280.2972 | 163.0388 | Coumaric acid |
| 107 | 4F | ID62_RP_neg | 49.3001 | 164.0641 | unknown |
| 108 | 4F | ID67_RP_neg | 47.5997 | 165.0402 | Arabinonic acid (98.25%) |
| 109 | 4F | ID72_RP_neg | 88.6861 | 167.021 | unknown |
| 110 | 4F | ID82_RP_neg | 280.4396 | 181.0507 | Hydroxyphenyllactic acid |
| 111 | 4F | ID85_RP_neg | 280.1192 | 182.0535 | unknown |
| 112 | 4F | ID97_RP_neg | 271.8839 | 195.0522 | Dimethyluric acid |
| 113 | 4F | ID98_RP_neg | 47.1844 | 195.0506 | Sugar acid |
| 114 | 4F | ID104_RP_neg | 346.4323 | 204.0671 | Indolelactic acid |
| 115 | 4F | ID107_RP_neg | 346.0053 | 205.0695 | unknown |
| 116 | 4F | ID136_RP_neg | 53.2571 | 259.0136 | 3-O-sulfo-β-D-galactose |
| 117 | 4F | ID147_RP_neg | 347.5474 | 272.054 | unknown |
| 118 | 4F | ID215_RP_neg | 44.3295 | 307.1504 | Fructoselysine |
| 119 | 4F | ID269_RP_neg | 570.8473 | 365.2306 | unknown |
| 120 | 4F | ID280_RP_neg | 46.2214 | 380.1617 | unknown |
| 121 | 4F | ID297_RP_neg | 492.7667 | 389.2698 | unknown |
| 122 | 4F | ID398_RP_neg | 492.7135 | 435.2752 | unknown |
| 123 | 4F | ID416_RP_neg | 39.8382 | 440.8856 | unknown |
| 124 | 4F | ID481_RP_neg | 58.5597 | 462.0931 | unknown |
| 125 | 4F | ID573_RP_neg | 418.5118 | 499.3269 | unknown |
| 126 | 4F | ID599_RP_neg | 429.2894 | 513.3428 | unknown |
| 127 | 4F | ID630_RP_neg | 53.4444 | 523.1444 | fucopyranosyllactose |
| 128 | 4F | ID656_RP_neg | 420.7276 | 536.3211 | unknown |
| 129 | 4F | ID677_RP_neg | 47.9845 | 546.2038 | N-acetyl-trisaccharide |
| 130 | 4B/4F | ID788_RP_neg | 282.616 | 811.4574 | unknown |
| 131 | 4F | ID8_HILIC_pos | 424.4444 | 60.0803 | unknown |
| 132 | 4F | ID80_HILIC_pos | 385.1537 | 123.0541 | 2-Acetylpyrazine |
| 133 | 4F | ID85_HILIC_pos | 351.9125 | 128.1056 | N-Acetylpiperidin |
| 134 | 4F | ID97_HILIC_pos | 369.6011 | 131.1162 | N-Acetylputrescine |
| 135 | 4F | ID137_HILIC_pos | 352.4391 | 145.1325 | N-acetylcadaverine |
| 136 | 4F | ID141_HILIC_pos<br>100% | 424.348 | 146.1171 | 3-Dehydroxycarnitine |
| 137 | 4F | ID145_HILIC_pos | 424.3643 | 147.1195 | Glutamine |
| 138 | 4F | ID161_HILIC_pos | 97.413 | 153.0638 | unknown |
| 139 | 4F | ID170_HILIC_pos | 332.2044 | 159.1481 | unknown |
| 140 | 4F | ID189_HILIC_pos | 405.4134 | 162.1117 | Carnitine |
| 141 | 4F | ID193_HILIC_pos | 405.2537 | 163.1126 | unknown |
| 142 | 4F | ID245_HILIC_pos | 312.4109 | 175.0703 | unknown |
| 143 | 4F | ID449_HILIC_pos | 439.5683 | 264.1192 | unknown |
| 144 | 4F | ID454_HILIC_pos | 439.7345 | 265.1215 | unknown |
| 145 | 4F | ID518_HILIC_pos | 357.7949 | 284.062 | unknown |
| 146 | 4B/4F | ID550_HILIC_pos | 45.7972 | 295.2258 | oxidized fatty acid |
| 147 | 4F | ID582_HILIC_pos | 44.1117 | 307.261 | stearic acid |
| 148 | 4F | ID607_HILIC_pos | 291.0532 | 314.231 | Decenoylcarnitine |
| 149 | 4F | ID614_HILIC_pos | 289.283 | 316.247 | Decanoylcarnitine |
| 150 | 4F | ID680_HILIC_pos | 284.8135 | 342.2629 | unknown |
| 151 | 4F | ID688_HILIC_pos | 283.692 | 344.2791 | Lauroylcarnitine |
| 152 | 4B/4F | ID768_HILIC_pos | 307.4853 | 371.218 | unknown |
| 153 | 4F | ID784_HILIC_pos | 459.039 | 381.1279 | unknown |
| 154 | 4F | ID795_HILIC_pos | 292.5264 | 384.2727 | Hydroxy-tetradecadiencarnitine |
| 155 | 4F/4H | ID807_HILIC_pos | 289.7417 | 386.2895 | Hydroxy-tetradecenoylcarnitine |
| 156 | 4F | ID839_HILIC_pos | 50.7572 | 397.378 | unknown |
| 157 | 4F | ID876_HILIC_pos | 416.8625 | 406.1316 | Lacto-N-biose I |
| 158 | 4F | ID899_HILIC_pos | 287.1428 | 412.3056 | Hydroxyhexadecadienoylcarnitine |
| 159 | 4F/4H | ID948_HILIC_pos | 43.7895 | 423.3233 | unknown |
| 160 | 4F | ID983_HILIC_pos | 274.8781 | 429.3742 | unknown |
| 161 | 4F | ID1001_HILIC_pos | 43.2323 | 431.3727 | unknown |
| 162 | 4F | ID1015_HILIC_pos | 395.6925 | 434.1883 | unknown |
| 163 | 4F | ID1021_HILIC_pos | 395.6508 | 435.1907 | unknown |
| 164 | 4F | ID1062_HILIC_pos | 42.4714 | 446.397 | unknown |
| 165 | 4F | ID1071_HILIC_pos | 42.6837 | 447.4 | unknown |
| 166 | 4F | ID1074_HILIC_pos | 42.8372 | 448.3988 | unknown |
| 167 | 4F | ID1105_HILIC_pos | 52.8675 | 459.3401 | unknown |
| 168 | 4F | ID1231_HILIC_pos | 461.7946 | 511.1628 | 3-Fucosyllactose |
| 169 | 4F | ID1345_HILIC_pos | 43.8975 | 548.5346 | unknown |
| 170 | 4F | ID1346_HILIC_pos | 97.4613 | 548.5369 | unknown |
| 171 | 4F | ID1380_HILIC_pos | 44.23 | 561.4887 | FAHFA |
| 172 | 4F | ID1399_HILIC_pos | 80.4379 | 567.4422 | unknown |
| 173 | 4F | ID1425_HILIC_pos | 46.2204 | 573.4763 | unknown |
| 174 | 4F | ID1431_HILIC_pos | 43.6126 | 576.566 | unknown |
| 175 | 4F | ID1565_HILIC_pos | 92.2118 | 630.6151 | unknown |
| 176 | 4F | ID1604_HILIC_pos | 227.4001 | 642.5123 | unknown |
| 177 | 4F | ID1621_HILIC_pos | 44.7479 | 650.6385 | Ceramide (d18:1/24:0) |
| 178 | 4F | ID1661_HILIC_pos | 40.811 | 663.4519 | unknown |
| 179 | 4F | ID1663_HILIC_pos | 41.092 | 664.4597 | unknown |
| 180 | 4F | ID1707_HILIC_pos | 45.8138 | 678.5186 | unknown |
| 181 | 4F | ID1726_HILIC_pos | 512.4292 | 689.2044 | unknown |
| 182 | 4F/4H | ID1780_HILIC_pos | 256.9355 | 706.5366 | PC(30:0) |
| 183 | 4F | ID1790_HILIC_pos | 98.0462 | 710.5901 | unknown |
| 184 | 4F/4H | ID1792_HILIC_pos | 97.5934 | 711.5925 | unknown |
| 185 | 4F | ID1794_HILIC_pos | 494.3389 | 714.242 | N-acetylated quadruple sugar |
| 186 | 4F | ID1812_HILIC_pos | 46.2407 | 719.57 | unknown |
| 187 | 4F | ID1822_HILIC_pos | 46.1982 | 721.5722 | unknown |
| 188 | 4F | ID1876_HILIC_pos | 44.335 | 749.601 | unknown |
| 189 | 4F | ID1989_HILIC_pos | 84.2318 | 792.6685 | unknown |
| 190 | 4F | ID1992_HILIC_pos | 86.5068 | 793.6716 | unknown |
| 191 | 4F | ID1994_HILIC_pos | 92.0972 | 794.6834 | unknown |
| 192 | 4F | ID1998_HILIC_pos | 76.3574 | 795.6861 | unknown |
| 193 | 4F | ID2074_HILIC_pos | 200.7976 | 819.6667 | unknown |
| 194 | 4F | ID2163_HILIC_pos | 42.3567 | 880.7645 | unknown |
| 195 | 4B/4H | ID990_HILIC_pos | 42.9535 | 430.3752 | unknown |
| 196 | 4H | ID989_HILIC_pos | 43.0046 | 429.3716 | unknown |
| 197 | 4H | ID971_HILIC_pos | 43.8123 | 427.3523 | unknown |
| 198 | 4H | ID924_RP_pos | 40.5003 | 320.8688 | unknown |
| 199 | 4H | ID580_RP_pos | 560.4512 | 263.2366 | unknown |
| 200 | 4H | ID503_HILIC_pos | 46.0612 | 279.2291 | Linolenic acid |
| 201 | 4H | ID2187_HILIC_pos | 548.1752 | 1022.3515 | unknown |
| 202 | 4H | ID2152_HILIC_pos | 530.8375 | 877.2995 | unknown |
| 203 | 4H | ID196_HILIC_pos | 44.9749 | 163.1463 | unknown |
| 204 | 4H | ID1912_HILIC_pos | 254.2377 | 763.599 | unknown |
| 205 | 4H | ID1906_HILIC_pos | 253.9488 | 762.5947 | PC34:0 |
| 206 | 4B/4H | ID1904_HILIC_pos | 253.4206 | 761.5871 | unknown |
| 207 | 4H | ID1889_HILIC_pos | 253.2523 | 758.5674 | PC(34:2) |
| 208 | 4H | ID1858_HILIC_pos | 256.0458 | 736.572 | unknown |
| 209 | 4H | ID1855_HILIC_pos | 255.7154 | 735.5715 | unknown |
| 210 | 4H | ID1852_HILIC_pos | 47.5158 | 734.5681 | PC(32:0) |
| 211 | 4H | ID1850_HILIC_pos | 253.782 | 734.5695 | PC(32:0) |
| 212 | 4B/4H | ID176_HILIC_pos | 427.8128 | 160.1324 | Aminovaleric acid betaine<br>(could be 4- or 5-) |
| 213 | 4H | ID1581_HILIC_pos | 44.1712 | 634.6276 | unknown |
| 214 | 4H | ID1501_RP_pos | 583.4205 | 428.3723 | Octadecanoyl-L-carnitine |
| 215 | 4H | ID1182_HILIC_pos | 471.4868 | 495.1841 | unknown |
| 216 | 4B/4H | ID1058_HILIC_pos | 42.9742 | 444.3705 | unknown |
| Fig4B | 4B | ID397_HILIC_pos | 357.6501 | 245.0794 | unknown |
| Fig4B | 4B | ID162_HILIC_pos | 45.5128 | 153.1254 | unknown |
| Fig4B | 4B | ID978_HILIC_pos | 43.2362 | 428.3824 | unknown |
| Fig4B | 4B | ID920_HILIC_pos | 60.6639 | 418.1883 | unknown |
| Fig4B | 4B | ID322_RP_neg | 516.766 | 399.1485 | unknown |
| Fig4B | 4B | ID2150_HILIC_pos | 529.651 | 876.2944 | unknown |

## STAR Methods

### Ethics

The study was carried out in accordance with the Declaration of Helsinki and was approved by the Ethics committee of the Technical University of Munich under the study number 254/17S and included the written informed consent of the parents or legal guardians of the participating infants. The study is registered at the German Clinical Trials Registry (DRKS00012313).

### Study participants

Healthy women in their third trimester of pregnancy were recruited in the greater Munich, Germany area and eligibility for participation in the study was assessed. Exclusion criteria were premature birth (gestational age being less than 37 weeks), physical deformities that affect feeding (such as cleft lip and palate) and serious medical conditions that required an intensive care stay exceeding 3 days. Based on current healthcare guidelines in Germany, parents were explicitly encouraged to breastfeed. For participation in the study, parents agreed to use the provided study infant formula in case formula feeding became necessary.

### Controlled intervention trial design

The Infantibio-II study was a double-blind, randomized and placebo-controlled interventional trial. Infant fecal samples were collected at age 0.5, 1, 3, 7 and 12 months with an additional, voluntary follow-up sample at age 2 years. Infants were randomized to one of four formula groups. The letters A-D were used for blinding of the different formula groups. Formula group A (placebo) contained no additional supplements. Formula group B contained two bacterial isolates from native fecal samples of infants^10^ belonging to the species *Bifidobacterium longum* subsp. *infanti*s and *B. breve*. Lactosan International GmbH & Co. used freeze-dried bacterial powder to supplement infant formula with 10^7^ cfu/g of each strain. Formula group C contained a syrup-based galacto-oligosaccharide (GOS) mixture (5g/L, FrieslandCampina). Formula group D contained both *Bifidobacterium* strains and GOS. Parents were free to decide if, when and how much formula was fed to the infants in addition to or after cessation of breastfeeding. The reference group were infants that were fully breastfed for the entirety of the study period with the exception of solid food in addition to breastmilk.

### Infant formula

Infant formula was manufactured by Töpfer GmbH (Dietmannsried, Germany) and provided blinded to the study team for distribution to participants. The four formulas could not be distinguished from one another by looks, smell or other characteristics. Unblinding of formula groups took place after study completion. The study formula was a powdered organic cows-milk based infant formula containing all the necessary nutrients based on laws and regulations at the time of manufacturing (2018-2020). Parents received detailed instructions on how to prepare drink-ready formula including a digital thermometer for correct preparation.

### Fecal sample collection

Parents collected fecal samples from the diapers of participating infants and recorded the time of sampling. Stool sampling kits were provided including collection tubes with DNA stabilizing solution (Invitek Moleculuar, 1038111300) and empty fecal collection tubes (Süsse Labortechnik, H8555T). Two spoons of fecal matter were added to the tubes and immediately refrigerated until pick-up by study team, latest 24 hours after defecation. Samples were aliquoted and stored at -80°C at the study center at the Technical University Munich in Freising, Germany (ZIEL-Institute for Food & Health). Case report forms detailing birth characteristics, familial health status, environmental factors and the current feeding habits of the infant were completed at enrollment of study and with every stool sample collection. Parental fecal samples (86 from mothers and 66 from fathers) were collected on a voluntary basis upon consent at the infant sample time point 1-month according to previously established protocols.^51^

### 16S rRNA gene amplicon sequencing and analysis

Metagenomic DNA was isolated from 600 µl fecal matter in DNA stabilizing solution using a previously described DNA-isolation protocol.^51^ Negative controls in form of DNA extracted from water and DNA extracted from stool stabilizer (without fecal matter) were included every 45 samples. DNA extraction was followed by two-step polymerase chain reaction (PCR) amplification and sequencing on Illumina HiSeq in paired-end mode (2x 250 bp) using the HiSeq Rapid Kit v2 from Illumina.^51^

The UPARSE-based^52^ platform “Integrated Microbial Next Generation Sequencing” (IMNGS)^53^ was adapted as an in-house offline version (NGSToolkit, v6.0.2-beta.5) for processing the sequencing output. Reads were trimmed to the first base. USEARCH 8.0^54^ was used for paring of samples with 1 allowed mismatch per barcode. Remaining reads were trimmed (5 nucelotides each) followed by denoising and clustering by 100% identity, creating zero-radius operational taxonomic units (zOTUS) for optimal resolution of microbial strains in 16S rRNA sequencing. Chimera sequences were removed using UCHIME.^55^ Following best practices, zOTUs with a relative abundance below 0.25% were removed.^56^ Taxonomic labels were assigned by SINA 1.6.1 with the SILVA database (release 128).^57^ The sequences of a subset of zOTUs of interest were identified using the EzBioCloud database.^58^ Maximum-likelihood phylogeny calculations were done with FastTree^59^ and visualized with EvolView.^60^ Longitudinal changes in microbial abundances were visualized using horizon plots generated with the BiomeHorizon R package (v1.0.0).^61^

RHEA was used for microbial profile analysis.^62^ Statistical analysis was performed in GraphPad Prism, version 9.4.1 (GraphPad Software, LLC) or in R (2022.02.3+492). Normalization (no random subsampling, no rounding; cutoff was set to 10000 reads for 1^st^ year samples, 20000 reads in case of the fermenter samples and infant samples including the 24-month samples. The different thresholds were chosen in relation to the number of reads of each dataset.) and analysis such as alpha- and beta-diversity as well as taxonomic binning were performed with Rhea. Correction for multiple testing was done with Benjamini-Hochberg procedure. For comparison between more than two groups, a one-way ANOVA was used for normally distributed data (Shapiro Wilk test, alpha = 0.05). Otherwise, we used a Kruskal-Wallis test with Dunn’s procedure as post-hoc multiple comparison test. Stars indicate significance level: *p < 0.05; **p <0.01; ***p < 0.001. The “NBClust (v3.0)” package in R was used for hierarchical clustering of samples. The optimal number of clusters was determined based on the Calinski and Harabasz index^63^ and Ward’s hierarchical clustering criterion.^64^ Differences in microbial composition were calculated using a PERMANOVA test on generalized UniFrac distances, where the length of the bar in the graph provides context for dissimilarities of samples.

PICRUSt2 (v2.4.2)^65^ was used for functional analysis on all zOTUs identified in infant samples. zOTU sequences were used for predicting Metacyc pathway abundances (Caspi, Billington et al. 2018) without considering super-classes. The estimated Metacyc pathway abundances were used as a basis for linear discriminant analysis (LDA) effect size (LEfSe) calculations (alpha: 0.01) comparing formula group A to the other feed groups.^66^

The bacterial transmissibility ratio (%) between mother and infant was determined based on the number of shared zOTUs present in the mother and the child divided by the total number of zOTUs present in the mother. A non-parametric Kruskal-Wallis test followed by a post-hoc Dunn test was used to assess the significance between the groups.

### Diurnal and circadian analysis of infant microbiota and *in vitro* samples

To identify diurnal rhythms, a cosine-wave equation with a fixed 24-h period: y = baseline + (amplitude*cos(2*π*((x-[phase shift)/24))) was fitted on alpha-diversity, taxa level and relative abundance of individual zOTUs. Significance was determined with an F-Test and goodness of fit was corrected for multiple testing. Significant rhythmicity was assumed to occur when p ≤ 0.05. An adjusted version of the “CompareRhythms” script^67^ created for circadian rhythm comparison between groups, which performs JTK_cycle^68^ followed by DODR.^69^ Heatmaps were created using the heatmap script in R. The amplitude shown in the Manhattan plot is based on JTK_cycle and the phase is calculated using cosine-wave regression. Circadian cosine-fitting for *in vitro* sample was performed using an in-house code by Isaiah Ting based on.^70^

### Shallow metagenomics sequencing

To further enhance the identification and the understanding of the role of individual bacterial strains, shallow shotgun metagenomic sequencing was performed using isolated gDNA on a NovaSeq machine (1G raw data with paired-end 150 bp) on a subset of paired infant samples at time points month 3 and month 7. The resulting metagenomic data was demultiplexed and from the raw sequencing reads adapters were trimmed using “Trim Galore” (developed by F. Krueger, The Babraham Institute: https://github.com/FelixKrueger/TrimGalore, v.0.6.7). Quality control of sequencing reads (removal of low-quality reads and contaminants) was done using the pipeline “Knead data” (https://github.com/biobakery/kneaddata, v.0.7.7-alpha). Species-level taxonomic annotation was done using MetaPhlan 3.0.13 and HUMAnN 3.0.^71^ Gene families that were detected via HUMAnN 3.0 were mapped to the KEGG pathway orthologs.^72^ Centered log ratio scaling was applied.

### Untargeted metabolomics

Sample preparation: 100 mg native infant fecal matter was added to a 15 ml tube (CKMix 50, Bertin Technologies) containing ceramic beads (mixture of 2.8 mm and 5.0 mm beads). Then, 5 ml of methanol-based dehydrocholic acid extraction solvent (c = 1.3 μmol/L) was added and the sample was homogenized using a bead beater system (Precellys Evolution, Bertin Technologies) with a cooling adapter containing liquid nitrogen (Bertin Technologies). The bead beating procedure was performed three times per sample for 20 seconds at 10 000 rpm with a 15 second break between each round of bead beating. The sample was then centrifuged at 8 000 rpm for 10 minutes at 10°C. 500 ml of clear supernatant was transferred to a LC-MS/MS vial for untargeted metabolomics analysis. Multiple smaller volumes of samples were pooled to prepare a QC-sample.

Untargeted mass spectrometric measurement: analysis was performed using a Nexera UHPLC system (Shimadzu, Duisburg, Germany) coupled to a Q-TOF mass spectrometer (TripleTOF 6600 AB Sciex, Darmstadt, Germany). Separation of the stool samples was performed either using a UPLC BEH Amide 2.1 × 100 mm, 1.7 µm analytic column (Waters, Eschborn, Germany) with a 400 µL/min flow rate or with a Kinetex XB18 2.1 x 100 mm, 1.7 µm (Phenomenex, Aschaffenburg, Germany) with a 300 µL/min flow rate. For the HILIC-separation the settings were as follows. The mobile phase was 5 mM ammonium acetate in water (eluent A) and 5 mM ammonium acetate in acetonitrile/water (95/5, v/v) (eluent B). The gradient profile was 100% B from 0 to 1.5 min, 60% B at 8 min and 20% B at 10 min to 11.5 min and 100% B at 12 to 15 min. For the reversed-phase separation eluent A was 0.1% formic acid and eluent B was 0.1% formic acid in acetonitrile. The gradient profile started with 0.2% B which was held for 0.5 min. Afterwards the concentration of eluent B was increased to 100% until 10 min which was held for 3.25 min. Afterward the column was equilibrated at starting conditions. A volume of 5 µL per sample was injected. The autosampler was cooled to 10 °C and the column oven heated to 40 °C. Every tenth run a quality control (QC) sample which was pooled from all samples was injected. The samples were measured in a randomized order and in the Information Dependent Acquisition (IDA) mode. MS settings in the positive mode were as follows: Gas 1 55, Gas 2 65, Curtain gas 35, Temperature 500 °C, Ion Spray Voltage 5500, declustering potential 80. The mass range of the TOF MS and MS/MS scans were 50–2000 m/z and the collision energy was ramped from 15–55 V. MS settings in the negative mode were as follows: Gas 1 55, Gas 2 65, Cur 35, Temperature 500 °C, Ion Spray Voltage –4500, declustering potential –80. The mass range of the TOF MS and MS/MS scans were 50– 2000 m/z and the collision energy was ramped from –15–55 V.

Data processing: The “msconvert” from ProteoWizard^73^ was used to convert raw files to mzXML (de-noised by centroid peaks). The bioconductor/R package xcms^74^ was used for data processing and feature identification. More specifically, the matched filter algorithm was used to identify peaks (full width at half maximum set to 7.5 seconds). Then the peaks were grouped into features using the “peak density” method.^74^ The area under the peak was integrated to represent the abundance of features. The retention time was adjusted based on the peak groups present in most samples. To annotate features with names of metabolites, the exact mass and MS2 fragmentation pattern of the measured features were compared to the records in HMBD^75^ and the public MS/MS spectra in MSDIAL,^76^ referred to as MS1 and MS2 annotation, respectively. Missing values were imputed with half of the limit of detection method, i.e., for every feature, the missing values were replaced with half of the minimal measured value of that feature in all measurements. To confirm a MS2 spectra is well annotated, we manually reviewed our MS2 fragmentation pattern and compared it with records in the public database, using SIRIUS^77^ or previously measured reference standards to evaluate the correctness of the annotation. The mass spectrometric data is made publically available using the massive repository (https://massive.ucsd.edu) under the accession number MassIVE MSV000093140 for the feces data and MassIVE MSV000093802 for the breastmilk and formula data.

### Metabolite analysis

To obtain information about potential associations between metabolites and species (data from shallow metagenomic sequencing) or zOTUs (data from 16S sequencing), network analysis was performed. For this, a correlation matrix was calculated. A spearman correlation was calculated and only features with a significant correlation and an absolute correlation value above 0.5 were selected. For plotting of the network, the R qgraph version 1.9 was used. Therefore payout spring was used, with a minimum absolut edge weight of 0.5.

Data from positive, negative and reverse phase were merged, i.e. some of the metabolites might appear more than once (due to being detected in different phases). To reduce the number of metabolites according to the post prevalent metabolites, the input tables were ordered by decreasing sum over all individuals. Afterwards a spearman correlation was calculated and used for further analysis. For 4F a subset of significantly different species was selected. Based on this subset a correlation matrix was calculated. In the heatmaps, only significant and high correlated (absolute correlation coefficient above 0.5) species - metabolites correlations are shown.

### *Ex vivo* Infant Colonic Continuous Fermentation

Infant Stool Samples Selection, Preparation, and Inoculation: Fecal samples of two 3-month old infants, one exclusively breast-fed and one exclusively fed with formula A, were selected to be representative of their feeding group based on Principal Component Analysis (PCA) and *Bifidobacterium* abundance. Fecal slurries were prepared under anaerobic conditions by diluting each sample (previously thawed on ice and weighted: BF = 1.03 g, A = 1.98 g) in reduced PBS (3% L-Cys HCl). Each slurry was filtered (70 µm cell strainer) and an equal amount was loaded in two syringes to be injected in duplicate chemostat vessels (BF = 2x6.5mL, A = 2x8mL). The whole procedure was finished ∼1.5 hours after the samples were thawed.

Preparation of the Culture Medium: A modified culture medium was prepared to mimic the physiological conditions of the infant colon. The media composition was based on the composition and calculations described in literature^78^ adapted to the composition of the infant formula A. A fat content of 10% was chosen for this experiment,^79^ and a full nutrient ratio of carbohydrate:nitrogen:fats after digestion with 13 g/L of carbohydrates was adjusted to 44:46:10 (corresponding to 11.88 g/L of fat). Therefore, the total concentration of undigested compounds in the medium were 2.96 g/L of total fat and 13.72 g/L of undigested N-compounds. The full composition of the culture medium, in g/L, was as follows (all the medium components were provided by Sigma-Aldrich, unless specified otherwise): *base media* (NaCl: 4.47; KCl: 4.43; MgSO_4_: 1.24; CaCl_2_: 0.1; NaHCO_3_: 1.5; KH_2_PO_4_: 0.5; FeSO_4_: 0.005; Bile salts: 0.05; Tryptone: 0.5; Peptone: 0.5; Yeast Extract: 2.5; Tween 80: 1; L-carnitine: 0.0015); *mucin media* (porcine gastric mucin type II: 4); and *milk media* (whey protein hydrolysate (Myprotein, UK): 11.21; casein: 0.47; pre infant formula A: 11.88). Each solution was sterilized by autoclaving at 121°C for 15 min (110°C for *milk media* after adjusting pH to 8.0^80^ and then mixed thoroughly. Two solutions were added after sterile filtration (0.2 μm): the first containing 0.8 g/L of L-cysteine, 5.86 g/L of lactose, and 0.004 g/L of inositol and the second was a vitamin solution as prepared by^81^ which also included hemin (0.01 g/L).

*Ex vivo* gut chemostat: The two infant fecal samples were cultured in duplicate chemostats (Bioreactor Model Multifors 2, Infors HT, Switzerland) according to the illustrated experimental scheme (**Fig. 5G**). In the first stage of the *ex vivo* gut chemostat model, a batch culture (total 48 h) was employed to allow for sample adaptation after inoculation. After 24 h without adjustment of pH or flow of fresh culture medium, the culture volume in the bioreactors were doubled and maintained at 800 mL until the end of the batch culture. In the second stage of the gut chemostat, a continuous culture was established for 7 days applying a retention time of 12.5 h.^78^ Samples were taken once each day at the same time (1.5 mL for DNA extraction and 16S rRNA Sequencing). Finally, the third stage were 48 hours of rhythmicity sampling during which samples were taken every 4 h. The following conditions were applied during the 9 days of continuous culturing: 400 mL of volume in the bioreactors, feed rate of 32 ml/h/chemostat, pH was set to 6.5 and then maintained (pH electrodes, Hamilton, Germany). Temperature was kept at 37°C, anaerobic conditions were controlled by supplying a continuous flow of N_2_ and redox values were measured (Redox electrodes, 1 per chemostat, Hamilton, Germany).

